# Stiffness of primordial germ cells is required for their extravasation in avian embryos

**DOI:** 10.1101/2022.05.09.491122

**Authors:** Daisuke Saito, Ryosuke Tadokoro, Arata Nagasaka, Daisuke Yoshino, Takayuki Teramoto, Kanta Mizumoto, Kenichi Funamoto, Hinako Kidokoro, Takaki Miyata, Koji Tamura, Yoshiko Takahashi

**Affiliations:** Frontier Research Institute for Interdisciplinary Sciences, Tohoku University, Sendai, Japan; Department of Ecological Developmental Adaptability Life Sciences, Graduate School of Life Sciences, Tohoku University, Sendai, Japan; Department of Biology, Faculty of Science, Kyushu University, Fukuoka, Japan; Department of Zoology, Graduate School of Science, Kyoto University, Kyoto, Japan; Department of Life Science, Okayama University of Science, Okayama, Japan; Division of Histology, Meikai University School of Dentistry, Sakado, Japan; Department of Anatomy and Cell Biology, Nagoya University Graduate School of Medicine, Nagoya, Aichi, Japan; Institute of Engineering, Tokyo University of Agriculture and Technology, Koganei, Japan; Institute of Fluid Science, Tohoku University, Sendai, Japan; Organization for Research Initiatives and Development, Doshisha University, Kyotanabe, Japan

## Abstract

During metastasis intravascularly circulating cancer cells undergo extravasation, which frequently takes place in vascular capillary beds ^1–3^. It remains poorly understood how the extravasation in the capillary beds is regulated. To address this question, chicken primordial germ cells (PGCs) serve as a powerful model since they circulate in blood stream and extravasate at a specific site of capillary bed near the forming gonad ^4–6^. The extravasation consists of two steps, the intravascular arrest of cells and their subsequent transmigration through the endothelial lining. We here demonstrate with live imaging at a single cell level *in vivo* that the arrest of PGCs is predominantly governed by occlusion at a narrow path in the capillary bed. In addition, this occlusion is enabled by a hightened stiffness of the PGCs, revealed by atomic force microscopy indentation analyses. The PGCs’ stiffness is regulated by actin polymerization: inhibition of the actin function causes not only a failure of PGC occlusion in the capillary bed, but also a failure of PGC colonization in the gonads at later stages. Following the occlusion, PGCs reset their stiffness to soften in order to squeeze through the endothelial lining as they transmigrate. The discovery of F-actin-mediated stiffness in pre-extravasating cells provides a model for understanding of dynamic mechanism by which other cells, including metastasizing cancer cells, extravasate in capillary beds.

---

PGCs in avian embryos exploit blood circulation to migrate from germinal crescent to the somatic gonad^5, 7^, during which time they need to undergo extravasation^4^. In early chicken embryos at Hamburger & Hamilton stage 10 (HH10)^8^, PGCs are widely distributed in blood vessels including extraembryonic vasculature. They are subsequently confined by HH16 to a specific region of the embryonic capillary bed (vascular plexus) near the pair of presumptive gonads, which are located posteriorly to the vitelline arteries (VA) running medio-laterally at the level of 20^th^ somite (Fig. 1 and Extended Data Fig. 1)^4^. The intravascularly trapped/arrested PGCs subsequently undergo transmigration to invade the splanchnic lateral plate (mesenteric primordium), and ultimately migrate to the gonad primordia^4, 9^. In this study, the region of vascular plexus accommodating extravasating PGCs is designated as Ex-VaP (<u>ex</u>travasation <u>va</u>scular <u>p</u>lexus). PGC-enriched Ex-VaP is confirmed by DDX4 for PGCs in chicken and also by double-labeling with DDX4 and QH1 (vasculature) in quail embryos (Extended Data Fig. 1).

**Figure 1.**
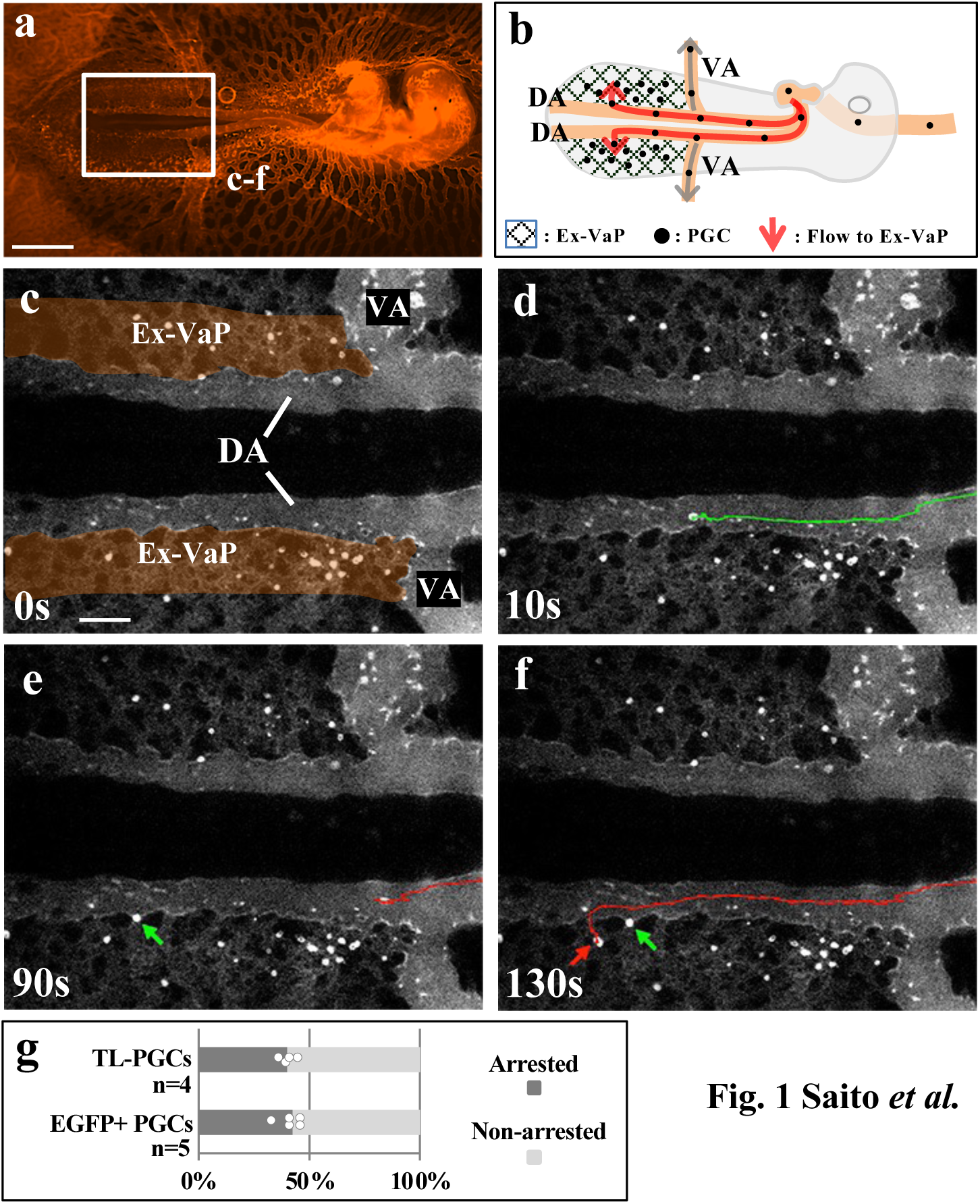
Circulating PGCs are arrested in Ex-VaP of HH15 chick embryo. **a,** Ventral view infused with fluorescent ink. **b,** Diagram of circulation and arrest of PGCs in Ex-VaP. **c-f,** Motion captures at 0s, 10s, 90s, 130s from the movie corresponding to the square in **a** (Supplementary video 1) show that some of TL-labeled endogenous PGCs (bright spots) circulating in the dorsal aortae are arrested in Ex-VaP (arrows). Green and red lines are tracks of two different PGCs. **g,** Quantification of arrested PGCs. TL-labeled endogenous PGCs (301 cells; 4 embryos), and back-infused EGFP+ PGCs (527 cells; 5 embryos) were assessed in Ex-VaP. DA, dorsal aorta; VA, vitelline artery. Scale bars: 1,000 μm in **a**, 200 μm in **c**.

A two dimensional (2-D) flat structure of early chicken embryos facilitates high speed recording of intravascularly labeled PGCs to visualize their arrest in Ex-VaP at a single cell level. At HH15, tomato lectin-conjugated FITC-labeled PGCs (TL-PGCs) in the pair of dorsal aortae (DA) move intermittently obeying the rhythm of heartbeat (Fig. 1 and Supplementary video 1). Whereas the majority of PGCs in DA turn into VAs at 20^th^ somite level and quickly flow out from embryo-proper, some PGCs keep circulating down posteriorly in DA, and finally turn laterally into Ex-VaP (Fig. 1 and Supplementary video 1). Since Ex-VaP is composed of meshwork-like narrow paths of vasculature, the velocity of PGC circulation is much lower than that in DA (150 μm/s and 300 μm/s in Ex-VaP and DA, respectively). Approximately 40% of the Ex-VaP-entering PGCs are arrested (Fig. 1 and Supplementary video 1), whereas the rest of the cells escape from the arrest and flow away toward extraembryonic vasculature, from which they probably circulate back into embryo-proper. The patterns of the circulation and arrest of TL-PGCs are comparable with those with CAGGS:EGFP-expressing PGCs (arresting rate: 40.9 ± 4.5%), which were stably transfected in *in vitro* by using transposon Tol2 system and back-transplanted/infused into host embryos (Fig. 1g, Extended Data Fig. 2 and Supplementary video 2)^10, 11^.

**Figure 2.**
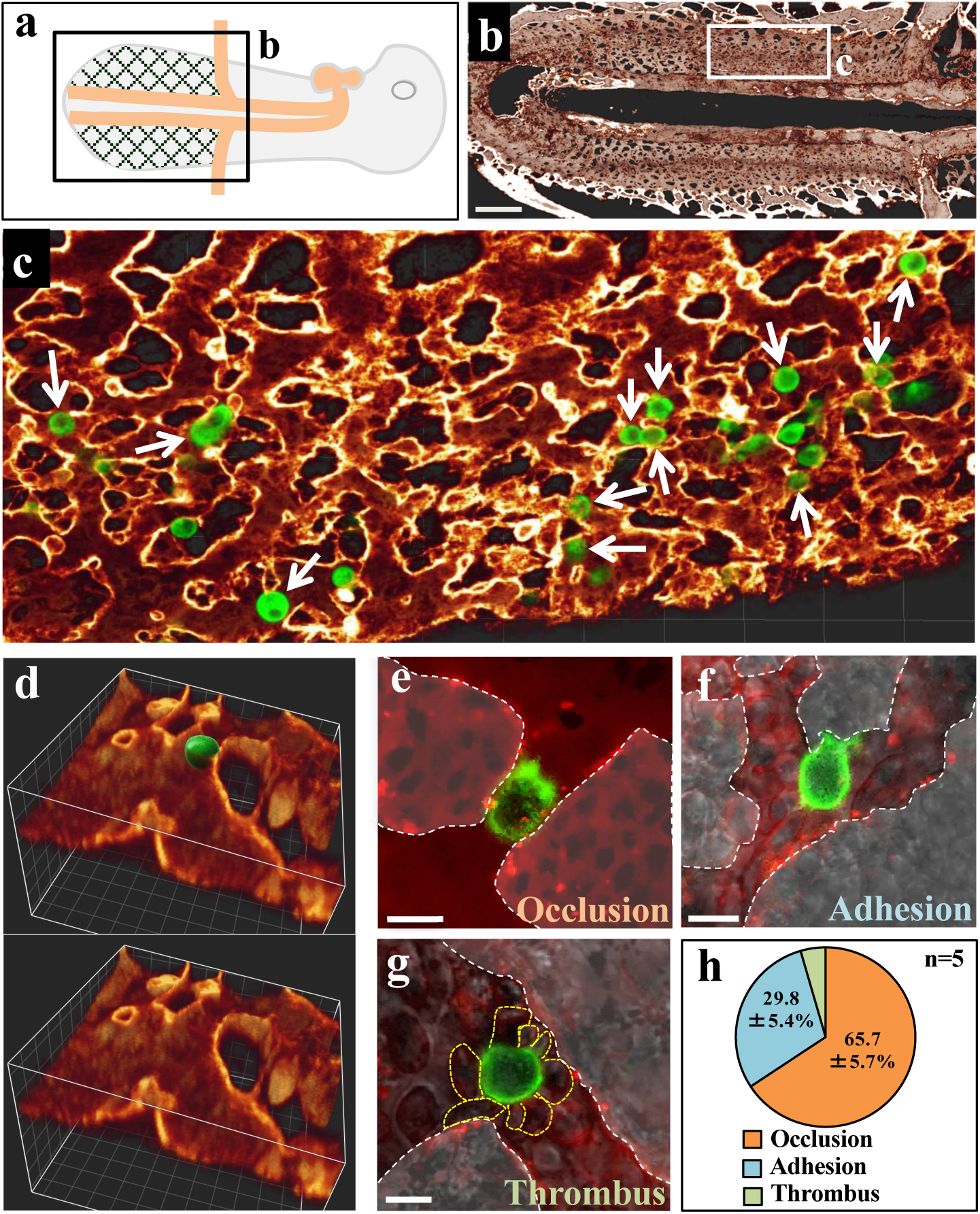
PGCs are occluded at constriction points in Ex-VaP. **a, b,** 3D reconstitution of QH1 signal in HH15 quail embryo corresponding to the square in **a. c,** Sliced and magnified view of the square in **b**. PGCs were labeled by DDX4 immunofluorescence (green). White arrows indicate occluded PGCs in Ex-VaP. **d,** 3D reconstitution image of occluded PGC and endothelial linings of Ex-VaP. PGC images were reprocessed to 3D iso-surface (green color with texture), and volume images of the endothelial linings (orange) were clipped with appropriate plane. **e-g,** Motion captures taken from Supplementary video 3 show occluded, adhered, and thrombus-like PGCs in Ex-VaP at high resolution. PGCs were labeled by TL- conjugated FITC. White and yellow dotted lines delineate the outline of endothelial linings and blood cells attached to the PGC, respectively **h,** Relative representation of three types of arrests in Ex-VaP, the occlusion, adhesion, and thrombus-like. Each value is a percentage to the total number of arrested PGCs in Ex-VaP of HH15 living embryos (n=5). Data were extracted from 15-minute movies after TL-conjugated FITC infusion. Scale bars: 300 μm in **b**, 10 μm in **e-g**.

A 3D-reconstructed image of vasculature at HH15 demonstrates a fine structure of DA through which PGCs enter Ex-VaP (Extended Data Fig. 3). In the ventro-lateral wall of DA, multiple passage branches (connection ports) connecting to Ex-VaP are observed, which are often associated with a pillar-like structure (Extended Data Fig. 3). Ports connecting to inter-somitic (intersegmental) vessels are also seen in the dorsal side of DA. The 3D-reconstructed vasculature has also revealed an intricate mesh-like structure of Ex-VaP that contains smallest (narrowest) paths of 10 μm wide (Fig. 2a-d and Extended Data Fig. 4), where PGCs are frequently occluded (65.7%; Fig. 2c-e, h and Extended Data Fig. 4). These occluded sites are designated as constriction points. In addition to the occluded PGCs, we have also found arrested PGCs that are not occluded at the constriction point.

**Figure 3.**
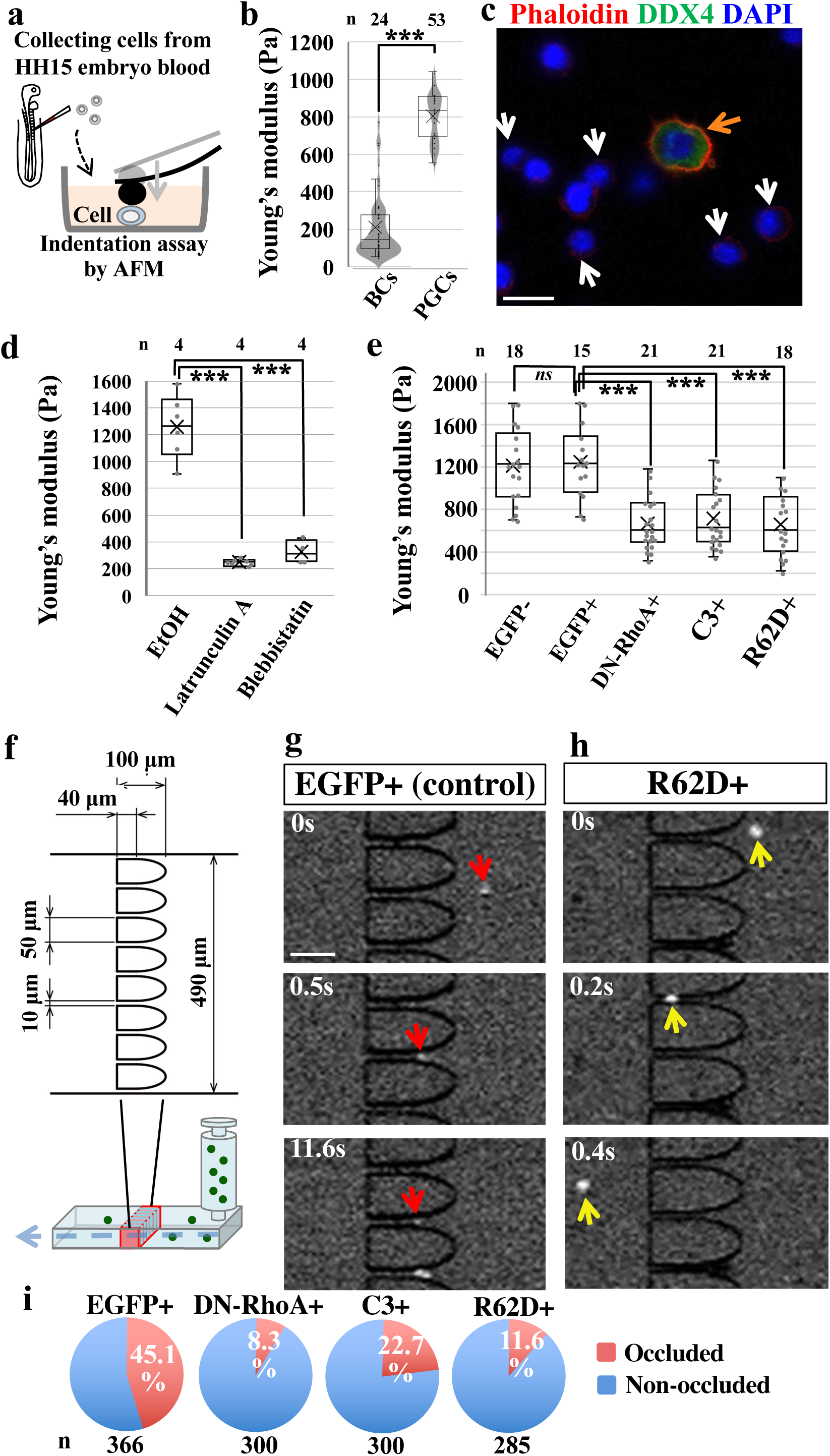
Actin-dependent stiffness of PGCs and occlusion assay. **a,** AFM indentation measurement applied to PGCs and blood cells (BCs) harvested from embryonic blood. Living cells attached to the dish bottom are pressed by a cantilever with a bead. **b,** Quantification of cell stiffness measured by AFM indentation tests. **c,** Immunofluorescence of DDX4 (green) and phalloidin staining (red) for PGCs (orange arrow) and BCs (white arrows). **d,** Cell stiffness of EtOH (control)- and chemical reagent-treated PGCs from HH15 chicken embryos. **e,** Cell stiffness of gene-manipulated PGCs. EGFP(+) and EGFP(-) are with and without Dox. **f,** Experimental design with a microfluidic device. **g, h,** Motion captures at indicated timepoints from the movies (Supplementary videos 4 and 7). A single PGC (red or white arrows) that flows from the right side passes through a slit after remaining at the slit for certain duration of time. **i,** Relative representation of occluded and non-occluded PGCs in the microfluidic constriction assay. Analyzed cell numbers are as indicated from 3 independent experiments. ns, not significant. ***p < 0.001. Scale bars: 10 μm in **c**, 50 μm in **g**.

**Figure 4.**
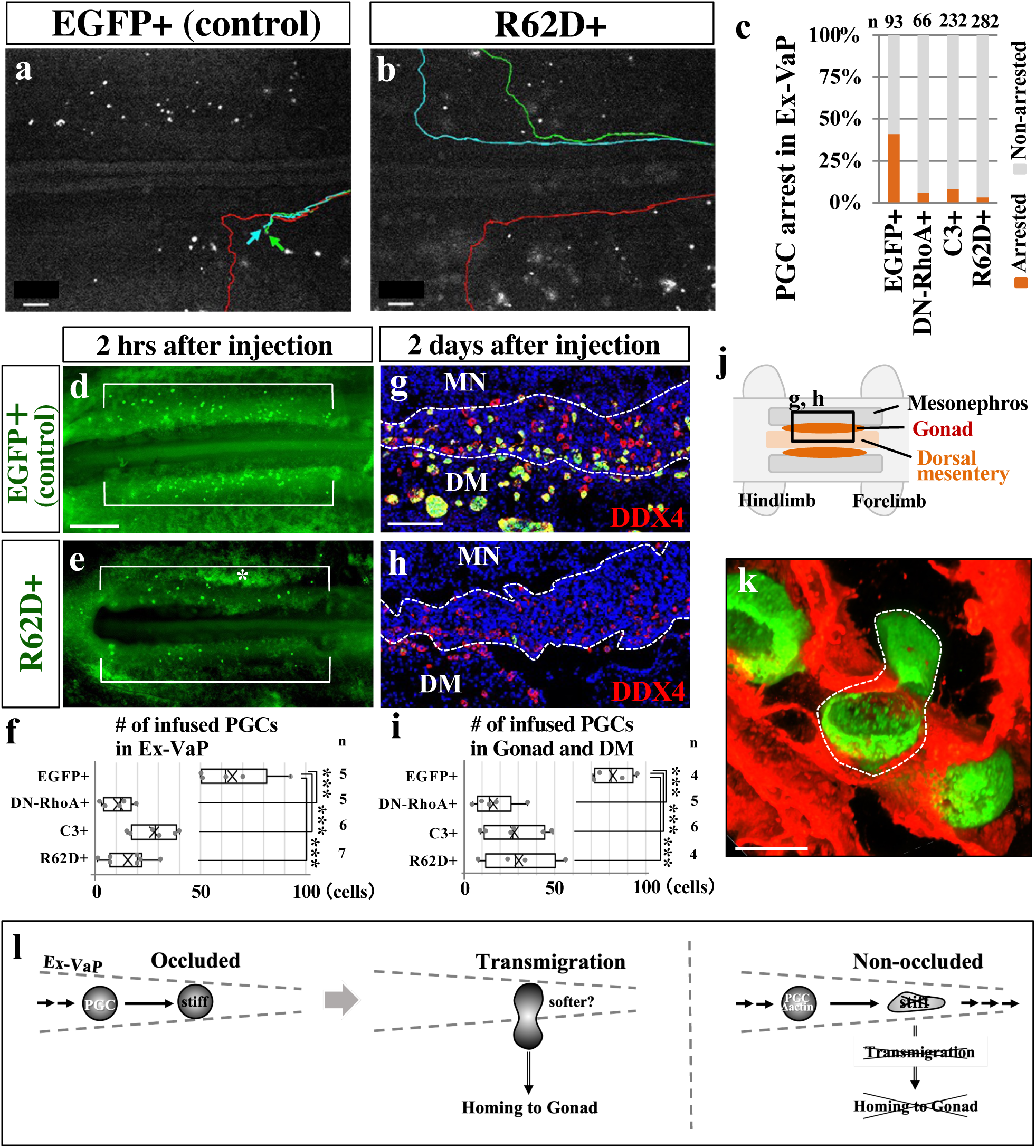
Actin-dependent stiffness of PGCs for the intravascular arrest, and its rapid reset for transmigration. **a, b,** Movie captures taken from Supplementary videos 8 and 11. Colored lines show tracks of intravascular translocations of PGCs in embryos. Arrows indicate an arrested PGC in Ex-VaP. **c,** Ratio of the arrest of actin-compromised PGCs in Ex-VaP. Analyzed cell numbers are as indicated from 5 embryos. **d, e,** Localization of back-infused PGCs (EGFP-labeled). EGFP immunofluorescence pictures in Ex-VaP (brackets) of HH15 embryos 2 hrs after infusion with EGFP+ or R62D+ PGCs. Asterisk shows noise signal. **f,** Quantification of arrested PGCs in Ex-VaP 2 hrs after infusion with 1,000 PGCs. **g, h,** DDX4 (red) and EGFP (green) immunofluorescence in horizontal sections of E4.5 chicken embryos (HH25) 2 days after infusion with EGFP- or R62D-transfected PGCs. Images correspond to a square in **j**. Dotted lines delineate the gonad primordium. MN, mesonephros; DM, dorsal mesentery. **i,** Quantification of infused PGCs in the gonads and DM. **j,** Ventral view of HH25 embryo. **k,** 3D reconstitution image of QH1 (red) and DDX4 (green) immunofluorescence of Ex-VaP in HH15 quail embryo. A squeezed PGC enclosed by a white dotted line in the course of transmigration through an inter-endothelial small hole in Ex-VaP. **l,** Summary diagram. Intravascularly circulating PGCs are highly stiff and successfully occluded at a constriction point in Ex-VaP. Subsequently, the occluded PGCs rapidly reset its stiffness to soften in order to squeeze themselves and transmigrate through an endothelial lining. The stiffness of PGCs is mediated by F-actin: F-actin-dysregulated PGCs are much softer, and fail to be occluded in Ex-VaP, resulting in poor colonization in the forming gonad. ***p<0.001. Scale bars: 200 μm in **a**, **b**, 500 μm in **d**, 100 μm in **g**, 10 μm in **k**.

To more precisely identify the distinguishing features of arrested PGCs in Ex-VaP, we have conducted high resolution live-imaging microscopy at the single cell level, and observed two more types of PGC’s arrest in addition to the occlusion type: one is an adhesion type in which PGCs are simply adhered to the endothelial lining regardless of the constriction point (29.8%) (Fig. 2f, h; Extended Data Fig. 4; Supplementary video 3).

The other type is a thrombus-mediated PGC arrest (4.5%), where single PGC is encapsuled by multiple blood cells (BCs; erythrocytes) forming an aggregate which is occluded as a whole (Fig. 2g, h; Supplementary video 3). In the following studies, we have scrutinized the mechanisms underlying the occlusion-mediated arrest of PGCs.

It is intriguing that whereas PGCs with a diameter of 13.1 ± 1.7 μm are occluded at the constriction point of 9-12 μm width, BCs of a comparable size, which are sporadically seen in the circulation, are not occluded (Fig. 2 and Extended Data Fig. 5). We have noticed that this difference is attributed to a deforming ability: PGCs retain their shape whereas the large BCs easily deform to pass through the constriction point (Fig. 2e and Supplementary video 3). These observations have raised the possibility that PGCs are highly stiff, enabling efficient occlusion in Ex-VaP.

To test this, the atomic force microscopy (AFM) indentation assay has been conducted to measure the stiffness of a single PGC prepared by harvesting the blood from HH15 chicken embryos (Fig. 3a)^12^. The elastic (Young’s) modulus obtained by vertical indentation on a single PGC is four times higher than that of a blood cell (Fig. 3b and Extended Data Fig. 6), indicating that PGCs are significantly stiff. For the AFM indentation assay, live PGCs are used, which are easily distinguished morphologically from other blood cells, and confirmed afterward by marker staining with DDX4 and Phalloidin (Fig. 3c, see also below).

Regarding factor(s) endowing PGCs with high stiffness, cytoskeletal components including actin fibers and microtubules are strong candidates^13–15^. Phalloidin staining reveals dense actin fibers distributed beneath the cortical membrane of PGCs, and the signal intensity in PGCs is five times higher than that in blood cells (Fig. 3c and Extended Data Fig. 7a). In contrast, α-tubulin amounts and its distribution patterns are comparable between PGCs and blood cells (Extended Data Fig. 7b).

To examine whether F-actin function is responsible for the PGC stiffness, actin inhibitors, latrunculin A (inhibiting actin polymerization) or blebbistatin (inhibiting acto- myosin function), have been added to cultured PGCs prepared from HH15 embryos. After 30 minutes, treated PGCs exhibit drastic reduction in cell stiffness revealed by the AFM assay (Fig. 3d and Extended Data Fig. 8). To further trace the F-actin-inhibited PGCs for a prolonged period of time, cultured PGC are gene-manipulated with dominant-negative type of RhoA (DN-RhoA)^16^, botulinum C3 enzyme (C3)^16^, or β-actin with a point mutation (R62D)^17^ known to act as a dominant negative effector against actin polymerization. Each of these genes is tet-on inducibly expressed in PGCs with Dox administration so that the actin inhibition commences upon the AFM assay but not during the *in vitro* culture for PGC propagation^18^. PGCs treated with each of these three inhibitors have reduced cell stiffness whereas EGFP-manipulation shows no effect (Fig. 3e). Thus, F-actin functions, but not those of microtubules, are necessary for endowing PGCs with stiffness. The cell diameter and the shape of PGCs are not affected by these inhibitors (Extended Data Figs. 5, 9).

To know whether the cell stiffness plays a major role in the occlusion of PGCs at a constriction point, we have prepared a microfluidic device with multiple micro-slits of 10 μm width and 50 μm height for each (Fig. 3f), which is equivalent with a narrowest width among vascular paths in Ex-VaP (Fig. 2 and Extended Data Fig. 4). PGCs transfected with tet-on inducible DN-RhoA, C3, or R62D are streamed into this device at flow velocity of 150-500 μm/s. The surface of the micro-slits has a low cell affinity due to hydrophobic property of PMDS, minimizing the effect on cell adhesion. In this assay, the occlusion is defined as follows: if cells remain at the micro-slit for longer than 5 seconds, they are regarded as occluded. In contrast, if cells pass through the slit within 5 seconds, they are regarded as non-occluded. Tet-on EGFP transfected-PGCs (control) are efficiently occluded (45.1%) (Fig. 3g, i and Supplementary video 4). In contrast, most of DN-RhoA, C3, or R62D transfected-PGCs are non-occluded (8.3%, 22.7%, 11.6%, respectively) (Fig. 3h, i, Extended Data Fig. 10 and Supplementary videos 5-7). These data indicate that the actin-mediated stiffness is essential for the PGC occlusion in a narrow path.

To determine whether the actin-mediated stiffness is also crucial for PGC occlusion/arrest in Ex-VaP *in vivo*, the aforementioned F-actin-inhibited PGCs (1,000 cells) are infused into host HH15 embryos. Control tet-on inducible EGFP-PGCs display normal patterns of intravascular circulation (arresting rate: 40.8% compared to Fig. 1g), arrests in Ex-VaP (65.2 ± 15.7 cells in Ex-VaP), and migration/homing to gonads (82.0 ± 9.5 cells/gonads) comparable with those for TL-labeled- and CAGGS:EGFP-PGCs (Fig. 4a, c, d, f, g, i, j, Extended Data Fig. 11 and Supplementary video 8), confirming that the tet-on system does not compromise normal behavior of PGCs *in vivo*. Dox is administered upon the infusion of the gene-manipulated PGCs so that the actin inhibition commences at this time point. Live-imaging analyses reveal that expression of DN-RhoA, C3, or R62D markedly reduces both the arresting rate (6.1%, 8.2%, 3.2%, respectively) (Fig. 4b, c, Extended Data Fig. 12 and Supplementary videos 9-11), and the total number of arrested PGCs in Ex-VaP (10.8 ± 6.2, 28.0 ± 9.3, 17.8 ± 8.5, respectively) (Fig. 4e, f and Extended Data Fig. 13). Importantly, the infused embryos show a significant failure of PGCs to migrate in the mesentery and to home gonads examined in E4.5 embryos (16.2 ± 10.2, 27.8 ± 14.6, 30.3 ± 17.4 cells in gonads, respectively) (Fig. 4h, i, j and Extended Data Fig. 11, 14). Collectively, the F-actin function-mediated stiffness is a crucial base for the PGCs to implement the efficient arrest in Ex-VaP, the prerequisite and essential step for the subsequent transmigration across the endothelial lining and migration/homing to gonads (Fig. 4l).

Following the PGC’s arrest in Ex-VaP, they complete the transmigration through the endothelial cell barrier in one hour (Fig. 4k). At this step, PGCs deform drastically in order to squeeze themselves to pass through an inter-endothelial space (hole), suggesting that the cell stiffness is quickly reset to soften the cell (Fig. 4l). Indeed, when polystyrene beads of 15 μm diameter are infused into a host embryo in a similar way to the PGC experiment, they are successfully arrested at the constriction points in Ex-VaP, but they never transmigrate (Extended Data Fig. 15)

We have demonstrated that the F-actin-mediated high stiffness is important for the PGCs to be efficiently occluded in Ex-VaP, a prerequisite step for their transmigration (Fig. 4l). Non-extravasating cells such as blood cells of a comparable size that are not occluded in Ex-VaP are much softer than PGCs. This unprecedented discovery of the requirement of cell stiffness for the extravasation highlights the possibility that other types of extravasating cells such as metastasizing cancer cells might also be highly stiff at the time of intravascular arrest prior to the transmigration. Indeed, the “mechanical trap theory” was proposed by James Ewing in 1928, which tried to explain the intravascular arrest of cancer cells^1, 19, 20^. However, since cancer cells *before* the intravascular circulation are not stiffer or even softer than normal cells^21, 22^, it remained unexplored whether these cells are/become stiffer at the time of their arrest in the capillary bed. It is increasingly appreciated that cancer cells and avian PGCs share common features regarding metabolism and gene expression profiles^23–28^. We have also shown that the stiffness of PGCs changes dynamically: following the occlusion, PGCs immediately reset the stiffness to soften so that they can squeeze themselves through an inter-endothelial small hole to transmigrate (Fig. 4k, l). Thus, intravascularly circulating cancer cells might also change their stiffness during different steps of extravasation. Investigations with a single cell level live-imaging using PGCs in avian embryos suggest new possible hypotheses to test concerning the mechanisms by which the extravasation of cancer cells is regulated.

## Acknowledgements

We thank Dr. Scott F. Gilbert for careful reading of the manuscript and discussion. We also thank Drs. T. Ogura and J. Kubo for the R62D construct. This work was supported by the following grants: JSPS KAKENHI (Grant number JP23116705, JP25111719, JP15K14358, 18H02445 for D. S., and JP20K21425, JP20H03259, JP 19H04775 for Y. T.) and Naito foundation, Japan health foundation, Narishige foundation, and Takeda foundation for D. S.

## Author Contributions

D. S. designed the study. D. S. and K. M. performed embryonic manipulations.

A. N., M. T. and D. S. performed and analyzed AFM experiments. R. T., H. K., T. T. and D. S. performed imaging experiments. D. Y. and K. F. designed microfluidic device, and D. S. performed the experiments and analyzed data. D. S., K. T. and Y. T. supervised the study. D. S. and Y. T. wrote the manuscript.

## Methods

### Animals, staging, and animal care

Fertilized chicken (*Gallus gallus domesticus*, Hypeco nera) eggs and fertilized quail (*Coturnix japonica*) eggs were purchased from Shiroyama poultry farm (Kanagawa, Japan) and from Nagoya University through the National Bio-Resource Project of the MEXT, Japan, respectively. Eggs were incubated at 38.5℃, and embryos were staged either by the somite number (described as “ss”) or by Hamburger and Hamilton’s stage^8^. All animal experiments were performed with the approval of the Institutional Animal Care and Use Committees at Tohoku University, Kyoto University and Kyushu University.

### Plasmid constructions

pT2A-CAGGS-EGFP: pT2AL200R150G vector^29^ was digested with XhoI-BglII. This site was blunt-ended, and inserted with the fragment of pCAGGS-EGFP containing the CMV enhancer, βActin promoter, EGFP, and polyA-additional sequences of the rabbit beta globin gene. pT2A-BI-TRE-EGFP: The pT2AL200R150G vector was digested with BglII-XhoI. This site was blunt-ended, and inserted with the fragment of pBI-Tight (Clontech) containing the bidirectional tetracycline-responsive element (TRE) with two minimal promoters of CMV in both directions, and two polyA-additional sequences of the rabbit beta globin gene. This vector was designated as pT2A-BI-TRE. The full-length of EGFP was amplified by PCR, and subcloned into the EcoRI-BglII site of pT2A-BI-TRE. pT2A-BI-TRE-EGFP- (DN-RhoA, C3, or βActinR62D (R62D)): The ORF of a dominant negative form of RhoA (DN-RhoA) or C3 transferase^16^ was subcloned into the MluI-EcoRV site of pT2A-BI-TRE-EGFP. The ORF of R62D (kind gift from Drs. Ogura and Kubo)^17^ was subcloned into the MluI-NheI site of pT2A-BI-TRE-EGFP. pT2A-CAGGS-Tet3G-2A-PuroR: pT2AL200R150G vector was digested with ApaI- BglII. PCR product including 2A peptide sequence flanking two multi-cloning sites and polyA-additional sequences of the rabbit beta globin gene was ligated into ApaI-BglII site of pT2AL200R150G. This vector was designated as pT2A-MCS1-2A-MCS2. CAGGS promoter sequence was inserted into SalI-PstI site of MCS1 of pT2A-MCS1- 2A-MCS2. Tet3G and PuroR PCR products were inserted into the NotI-SphI site of MCS1 and MluI-NheI site of MCS2, respectively.

### Establishment and maintenance of PGCs in culture

Circulating PGCs along with blood cells were harvested from blood of HH 14 chicken embryos, and were cultured in calcium-free DMEM (Gibco) diluted with water, containing <u>F</u>GF2 (Wako), <u>A</u>ctivin A (Peprotech), and <u>c</u>hicken <u>s</u>erum (FAcs medium) according to the method previously described^11, 30^. After one month, expanded PGCs were cryo-preserved at -80℃ in Bambanker (NIPPON Genetics) until used for experiments.

### Plasmid transfection and establishment of gene-manipulated PGC lines

2 x 10^4^ cultured PGCs were washed with OPTI-MEM (Gibco). PGCs were suspended in 10 μl of electroporation re-suspension buffer (R buffer) (Invitrogen). After addition of plasmid DNA (1 μg), they were electroporated by Neon Transfection System (Invitrogen) with optimized condition (1,300 V, 10 ms pulse width, 3 pulses). We used 5 different sets of plasmids (pT2A-CAGGS-EGFP + pCAGGS-T2TP^10^; pT2A-BI-TRE-EGFP + pT2A-CAGGS-Tet3G-2A-PuroR^16^ + pCAGGS-T2TP; pT2A-BI-TRE-EGFP- DN-RhoA + pT2A-CAGGS-Tet3G-2A-PuroR + pCAGGS-T2TP; pT2A-BI-TRE-EGFP- C3 + pT2A-CAGGS-Tet3G-2A-PuroR + pCAGGS-T2TP; pT2A-BI-TRE-EGFP-R62D + pT2A-CAGGS-Tet3G-2A-PuroR + pCAGGS-T2TP). Transfected PGCs were seeded into antibiotics-free FAcs medium. After 12 hours, the medium was exchanged to the conventional FAcs medium.

Following pT2A-CAGGS-EGFP transfection, each PGC was separately seeded into 96-well plate dish to obtain EGFP-positive colonies. PGCs receiving the PuroR gene were cultured in the FAcs medium containing 0.5 μg/ml puromycin for two weeks to enrich puromycin-resistant cells.

### Immunofluorescence staining and phalloidin staining

For DDX4 and α-Tubulin immunostaining and phalloidin staining in floating PGCs, we mounted living cells with Smear Gell^TM^ (GenoStaff) on an APS-coated slide glass according to the manufacture’s instruction. Cells in the slide glass were fixed with 4% PFA/PBS (paraformaldehyde/phosphate buffered saline) for 30 minutes at room temperature (RT). Specimens were washed in PBS and blocked with 1% Blocking reagents (Roche)/TNT (0.1 M Tris-HCl (pH 7.5), 0.15 M NaCl, 0.1% Tween 20) for one hour at RT. The slide glasses were incubated at 4℃ overnight with anti-DDX4 rabbit polyclonal antibody (custom-made by Eurofins genomics, 1:10,000), or anti-α-Tubulin antibody (mouse T6199, SIGMA, 1:4,000) in the blocking solution. After 3 washes in TNT for 5 minutes each, the specimens were reacted with anti-rabbit IgG-Alexa 488- conjugated goat antibody (Invitrogen) diluted 1:1,000 or anti-mouse IgG-Alexa 488- conjugated goat antibody with DAPI, Alexa Fluor 647 Phalloidin (ThermoFisher, 1:100), and blocking solution for one hour at RT. The specimens were washed 3 times in TNT and sealed with FluoSave reagent (Calbiochem).

Immunostainings with DDX4 (1:2000), QH1 (mouse, Hybridoma Bank, 1:2) and EGFP (goat, abcam, 1:1000) in whole embryos, dissected gonad/mesonephros, or histological sections were performed as previously described^7, 31, 32^. Frozen sections of fixed chicken embryos (10 μm thick) were prepared with a cryostat (Microm, HM500 OM).

### AFM indentation measurement and chemical treatments

To attach PGCs or blood cells on a plastic dish, dishes were pre-coated with 0.01% poly-L-lysine solution (SIGMA) or 0.1% poly-ethylenimine (SIGMA) for 30 minutes at 4℃. After 3 times washes in PBS, cells were seeded in a dish with Dulbecco’s modified Eagle medium/Ham’s F-12 medium (SIGMA), and incubated for 30 minutes for 37℃ in 5% CO2. After adding Latrunculin A (Cayman Chemical) or blebbistatin (Toronto Research Chemicals), floating cells were discarded.

All measurements (quantification) were performed using Cellhision200 (JPK Instruments, Berlin, Germany) mounted on an IX71 inverted microscope (Olympus). The equipment for indentation measurement and determination of the spring constant of cantilevers were previously shown^12^. The cantilever was equipped with a borosilicate bead (sQUBE, CP-CONT-BSG, 10 μm diameter). The applied force was 1 nN, and approach and retraction velocities were 650 nm/s. Each measurement point was set at the top of a cell. Force-distance curves were prepared with JPK DP software v.5 (JPK Instruments). In all cases, the indentation depth was 1.0 μm, and the spring constant of cantilever was determined prior to the measurements using the thermal noise method in air (nominal value, 0.2 N/m).

### TL-injection, PGC- and bead infusion, Dox administration, and embryo culture

Tomato lectin (TL)-conjugated FITC (SIGMA) was diluted by OPTI-MEM (50 μg/ml) for PGC visualization. To visualize both PGCs and blood plasma, CellMask^TM^ orange plasma membrane stains (Life Technologies) was added to the TL-FITC working solution at a 1:1,000 dilution. 1 μl of TL-FITC with/without CellMask was injected into the heart of HH15 embryos by a fine glass capillary (Narishige, GD-1) prepared with the puller (Narishige, PC-10).

For back-infusion, cultured PGCs were collected by centrifugation at 100 g for 5 minutes with several washes in OPTI-MEM. The number of living cells was adjusted to 1,000 cells/μl, and 1 μl of such suspension was injected into the heart of HH15 embryos by a fine glass capillary. For the tet-on induction, 1 μg/ml Doxycycline (Dox) (Clontech) was added into PGC-cultured FAcs medium 12 hours before the injection. Red fluorescent beads of 15 μm diameter (Invitrogen, FluoSpheres^TM^ Polystyrene Microspheres) were washed three times in PBS, and suspended in OPTI-MEM. The number of beads was adjusted to 1,000 beads/μl, and 1 μl of such suspension was infused into the heart of HH15 embryos with a glass capillary. Dox administration *in ovo* was as previously described^18^. For live-imaging, manipulated embryos were *ex-vivo* cultured as shown previously^33^.

### Microfluidic devices for cell arrest assay

In fabrication of the microfluidic device, the channel pattern with nine 10 µm- slits was transferred from silicon wafer to polydimethylsiloxane mold (PDMS; Silgard 184 Silicone Elastomer Kit, Dow Corning, USA) by soft lithography. Holes for the inlet and outlet with the diameter of 2 mm were punched to access the channel. The patterned side of the PDMS mold and a glass cover slip were bonded with each other after plasma treatment for 1 min and 40 seconds. The channel was filled with sterilized ultrapure water for preservation. In order to provide hydrophobic property to the device, we dried the device at 60℃ for 24 hours prior to the experiments.

Dox treated-PGCs suspended in OPTI-MEM (1,000 cells/μl) were loaded in 1ml syringe (Terumo), and pumped to the microfluidic device at 7 pl/min by syringe pump Legato 110 (KD Scientific).

### Capture and process of images

For live imaging, cultured embryos or microfluidic device were set in a humid chamber at 37 ℃ . Movies were taken by a cooled CCD camera, ORCA-R2 (HAMAMATSU Photonics) attached to the macro zoom microscope MVX10 (Olympus) with the software High Speed Recording (HAMAMATSU Photonics), or A1R confocal microscope (Nikon). Obtained images were processed with Manual Tracking Tool of ImageJ (https://imagej.nih.gov/ij/). For fixed cells and tissue sections, images were obtained with SP5 confocal microscope (Leica). Images of fixed whole samples were obtained with M205 FA stereomicroscope (Leica) or A1R confocal microscope (Nikon). Acquired Z-series images were deconvoluted and processed for 3D reconstruction by using Huygens (Scientific Volume Imaging) and Imaris software (ver.9.5, Oxford Instruments), respectively. Regions of interest (ROIs) showing PGC occlusion in narrow vascular plexus were selected from the whole image. The PGC images were further reconstructed to 3D iso-surfaces with texture, and volume images of plexus structure were clipped with appropriate x-y plane(s).

### Statistical analysis

All box plots represent the mean, upper and lower interquartile, error bars (s.e.m) with median (x). P values were obtained by a 2-tailed, unpaired Student’s t test (Excel). Box plots and bar graphs were made by Excel, and a violin plot was made by RAWGraphs.

**Supplementary Video 1**

**Intravascular circulation of endogenous PGCs and their arrest in Ex-VaP.** The movie was recorded at the posterior portion of HH15 chicken embryo by ORCA-R2 attached to MVX10 at 10 frames/s, and played at the normal time rate (10 frames/s). PGCs was labeled by TL-FITC. Motions of two PGCs were traced by green and red lines, and the arrested position were pointed by green and red arrows. Anterior is to the right.

**Supplementary Video 2**

**Intravascular circulation of back-infused PGCs and their arrest in Ex-VaP.** *In vivo* time-lapse movie of CAGGS:EGFP-expressing PGCs. PGCs were infused 15 min before the recording. See legends of Supplementary Video 1. Motions of four representative PGCs were traced by green, red, light blue, and purple lines, and their arrested positions were pointed by arrows.

**Supplementary Video 3**

**Microscopic views of endogenous PGCs’ arrest in Ex-VaP in HH15 chicken embryo.** The movies were recorded by A1R confocal microscope at 8 frames/s, and played at the same time rate. PGCs were labeled by TL-FITC. Blood plasma, endothelial cell walls, and BCs were visualized by CellMask^TM^ orange plasma membrane stains. Three modes of PGC arrest (occlusion, adhesion, and thrombus-like) were shown.

**Supplementary Video 4**

**Constriction assay with EGFP+ PGCs (control) by microfluidic device.** The movie was recorded by ORCA-R2 attached to MVX10 at 10 frames/s, and played at the same time rate. EGFP+ PGCs were streamed by micropump at 150-500 μm/s.

**Supplementary Video 5**

**Constriction assay with DN-RnoA+ PGCs by microfluidic device.** See legends of Supplementary Video 4.

**Supplementary Video 6**

**Constriction assay with C3+ PGCs by microfluidic device.** See legends of Supplementary Video 4.

**Supplementary Video 7**

**Constriction assay with R62D+PGCs by microfluidic device.** See legends of Supplementary Video 4.

**Supplementary Video 8**

**Intravascular circulation of back-infused EGFP+ PGCs (control) and their arrest in Ex-VaP.** The movie was recorded by ORCA-R2 attached to MVX10 at 10 frames/s, and played at 4x speed. EGFP+ PGCs were infused 15 min before the recording. Dox was administrated 12 hrs before the infusion. Motions of three representative PGCs were traced by green, red, and light blue lines, and their arrested position were pointed by arrows.

**Supplementary Video 9**

**Intravascular circulation of back-infused DN-RhoA+ PGCs and their arrest in Ex-VaP.** See legends of Supplementary Video 8.

**Supplementary Video 10**

**Intravascular circulation of back-infused C3+ PGCs and their arrest in Ex-VaP.** See legends of Supplementary Video 8.

**Supplementary Video 11**

**Intravascular circulation of back-infused R62D+ PGCs and their arrest in Ex-VaP.** See legends of Supplementary Video 8.

---

**Extended Data Figure 1.**
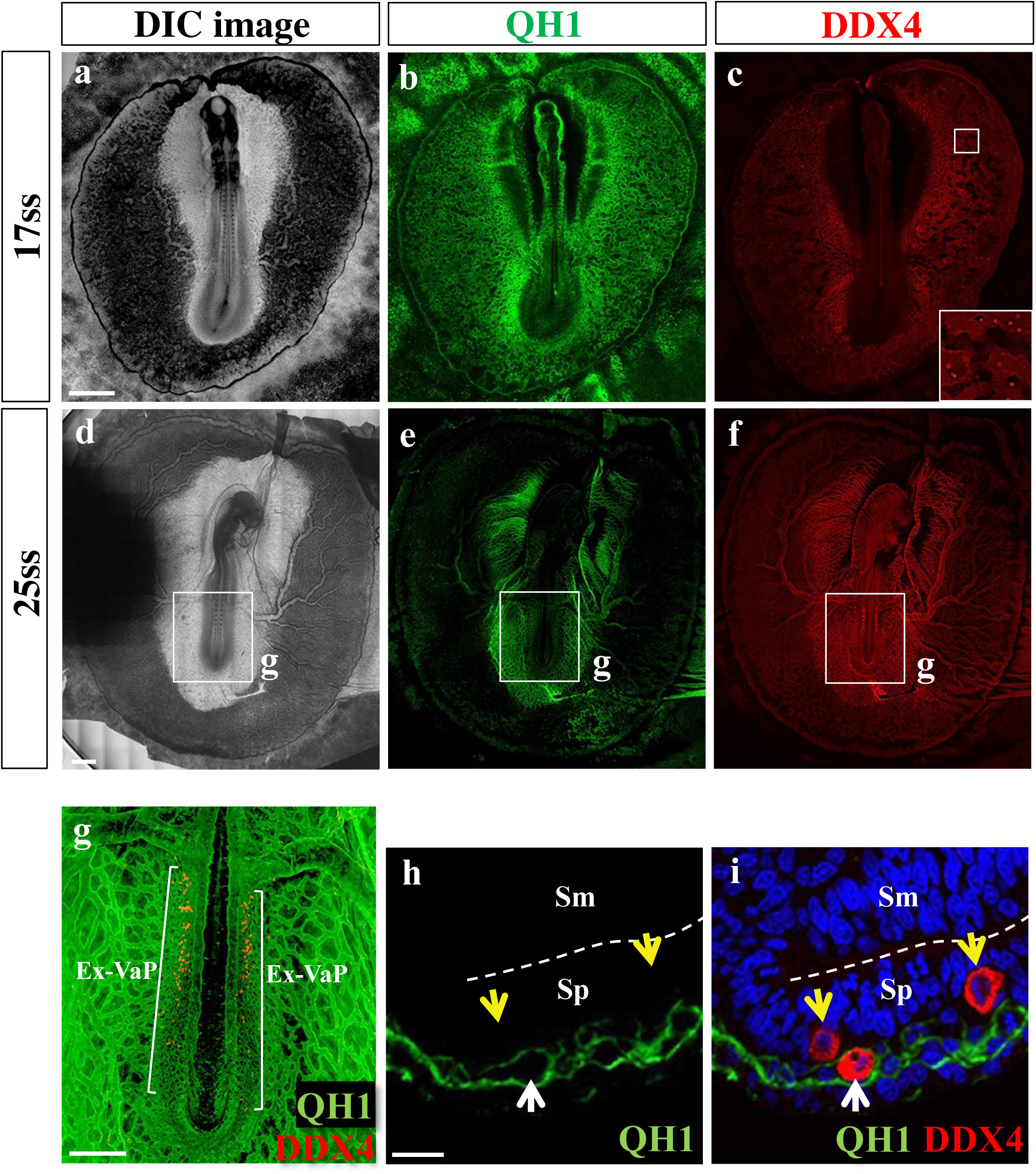
Intravascular PGCs accumulate and extravasate in Ex-VaP. a-f, QH1 (green) and DDX4 (red) immunofluorescence and DIC images of 17 somite stage (ss) (HH12, **a-c**) and 25ss (HH15, **d-f**) quail embryos. The inset in **c** is a magnified view of the square. **g,** Magnified and merged image of the square in **d-f**. White brackets indicate the Ex-VaP area. **h, i,** QH1 and DDX4 immunofluorescence and DAPI (blue) signals in the Ex-VaP transverse section of 27ss (HH15) quail embryos. Yellow and white arrows indicate extravascular and intravascular PGCs, respectively. Dotted lines delineate coelomic cavity. Sm, somatopleure. Sp, splanchnopleure. Scale bars: 1000 μm in **a**, **d,** 500 μm in **g**, 20 μm in **h**.

**Extended Data Figure 2.**
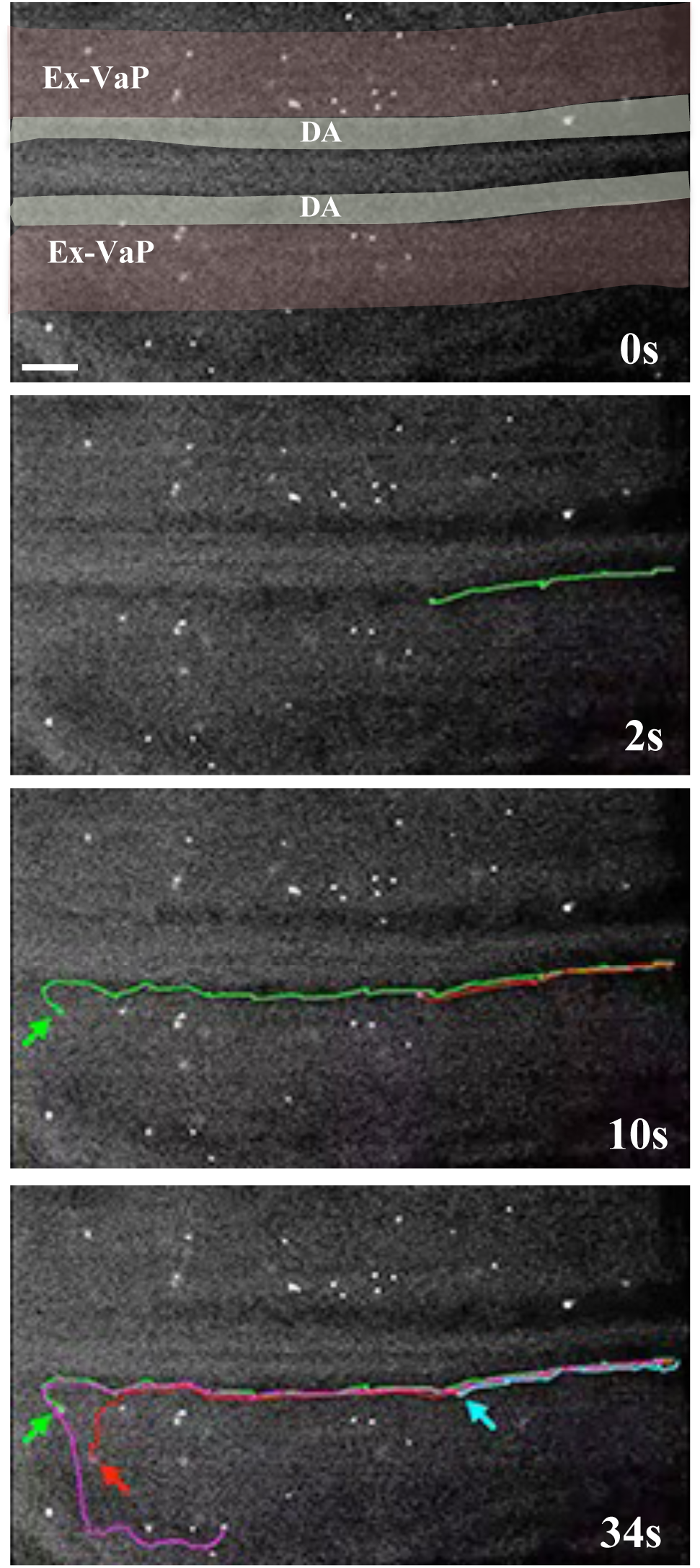
Back-transplanted/infused PGCs are preferentially arrested in Ex-VaP. Serial motion captures from Supplementary video 2 showing infused EGFP+ PGCs arrested in Ex-VaP (arrows) of HH15 chicken embryo. Green, light blue, and red lines depict tracks of PGCs’ translocation. Scale bar: 200 μm.

**Extended Data Figure 3.**
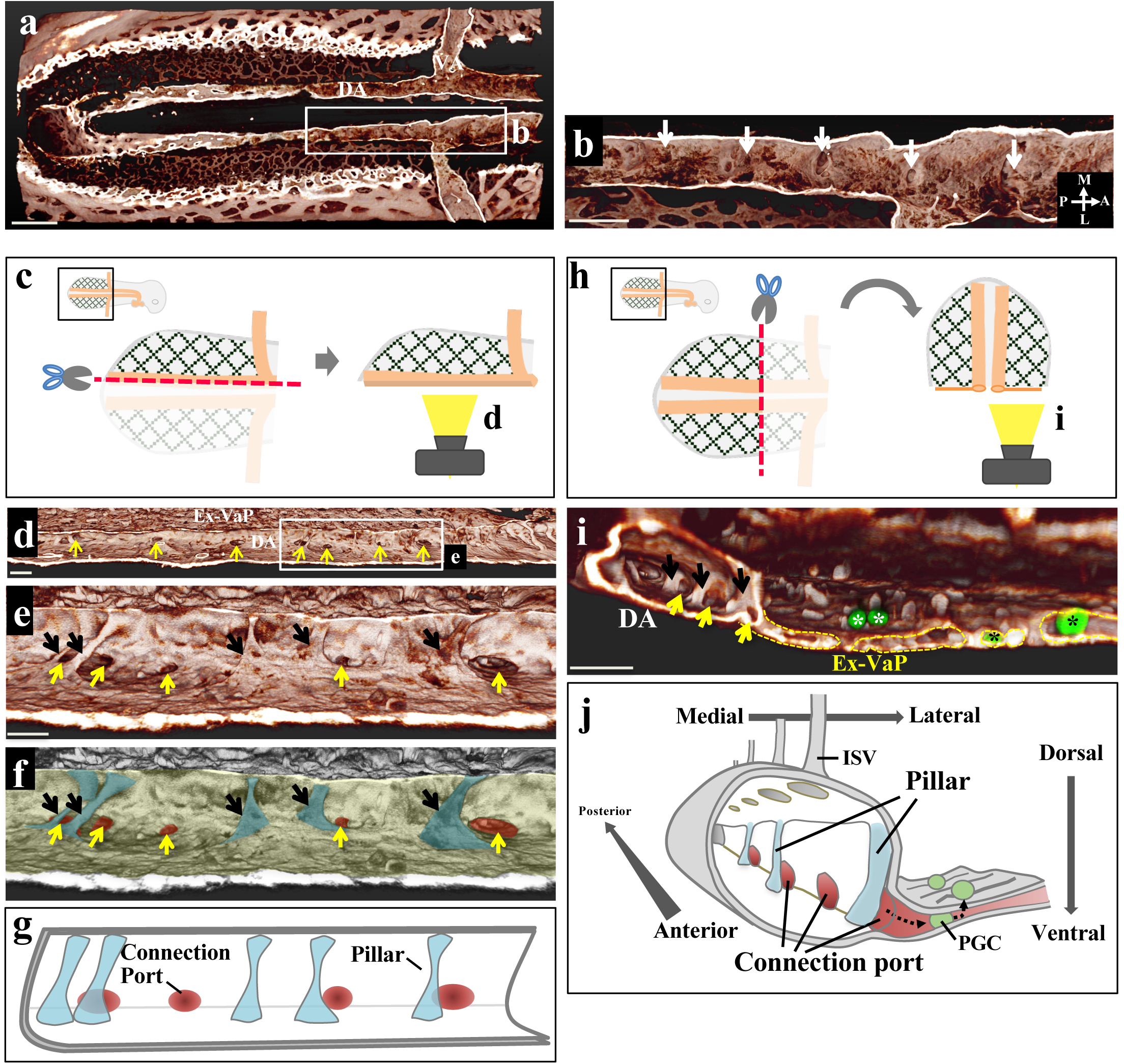
3D structures of dorsal aorta of HH15 quail embryo. **a,** 3D reconstructed image of QH1 signals in a posterior region including DA and Ex-VaP. The image is horizontally clipped and viewed from the ventral side. **b,** Magnified view of the square in **a**. White arrows indicate ports in the dorsal wall of DA connecting to intersegmental vessels. **c,** Diagram explaining views of **d-f** images. 3D reconstructed image of QH1 signals is sagittally clipped along the left DA, and viewed from the lateral right side. **d,** View as explained in **c**. Yellow arrows indicate the ports connecting to Ex-VaP (connection ports). **e,** Magnified view of the square in **d**. Black arrows show intravascular pillars near the connection ports in DA. **f,** The connection ports, pillars, and DA wall are highlighted in different colors. **g,** A trace diagram of **f**. **h,** Diagram explaining the view of **i** image. 3D reconstructed image of QH1 signals is transversely clipped and viewed from the anterior side. **i,** Anterior view of the left DA and Ex-VaP. Connection ports (yellow arrows) opening to Ex-VaP and pillars (black arrows) are seen. Yellow dotted lines delineate transverse surface of Ex-VaP. PGCs are immuno-stained by DDX4 antibody (green). Black and white asterisks indicate PGCs inside and outside of Ex-VaP, respectively. **j,** Topographical diagram showing DA, connection ports, pillars, Ex-VaP, and PGCs. Scale bars: 300 μm in **a**, 150 μm in **b**, 70 μm in **d,** 50 μm in **e**, 100 μm in **i**.

**Extended Data Figure 4.**
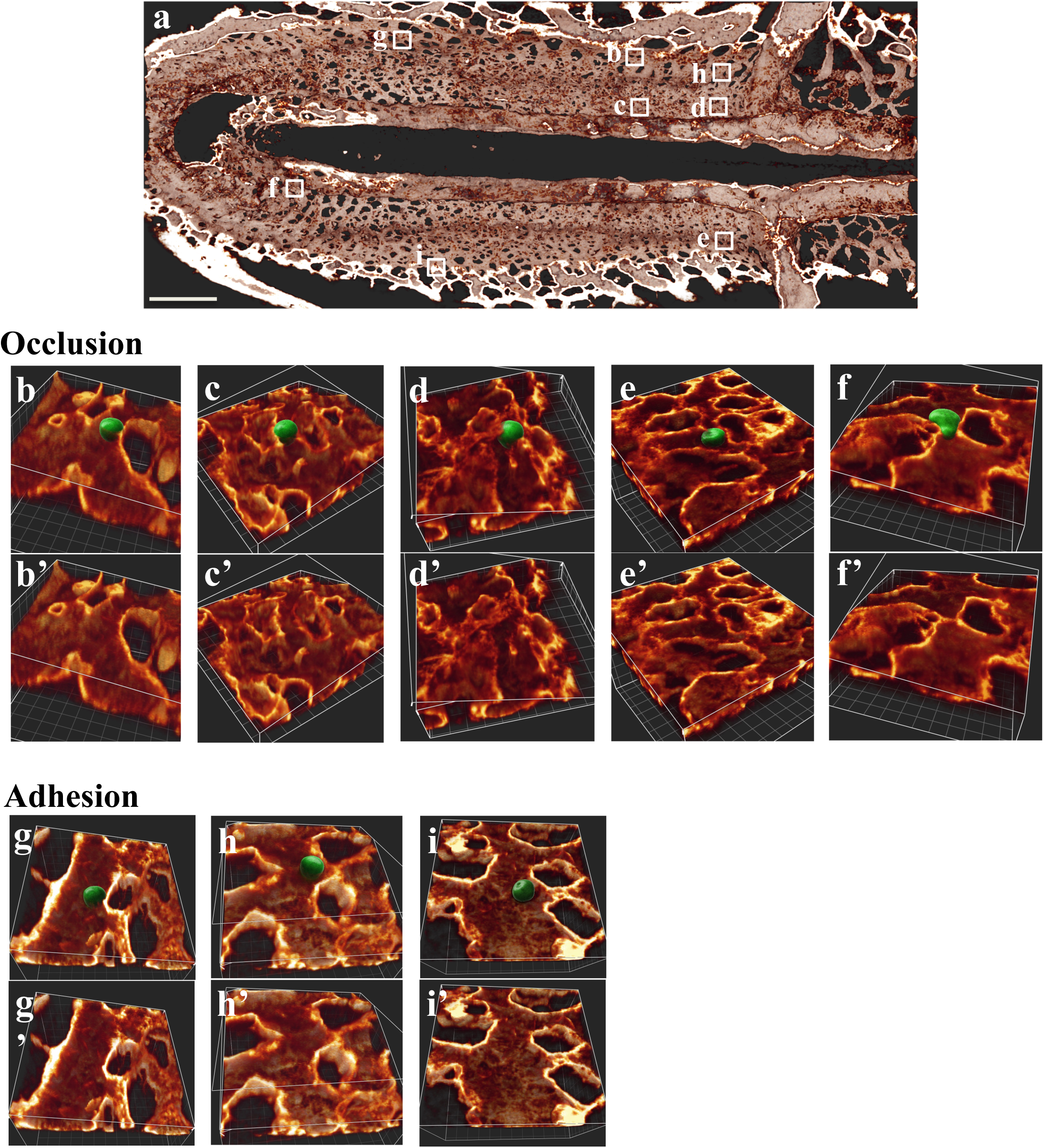
Occluded and adhered PGCs in Ex-VaP. **a,** 3D reconstructed image of QH1 signals in HH15 quail embryo. The image is clipped and viewed from the dorsal side. Scale bar: 300 μm. **b-f,** PGCs (DDX4+, green) are occluded in constriction points in Ex-VaP. **g-i,** PGCs adhere to an Ex-VaP endothelial wall. PGC images are reprocessed to 3D iso-surface (green color with texture), and then volume images of the endothelial wall (orange) are clipped with appropriate plane to facilitate viewing of PGCs and endothelial structure. Each image is obtained from the square region in **a**.

**Extended Data Figure 5.**
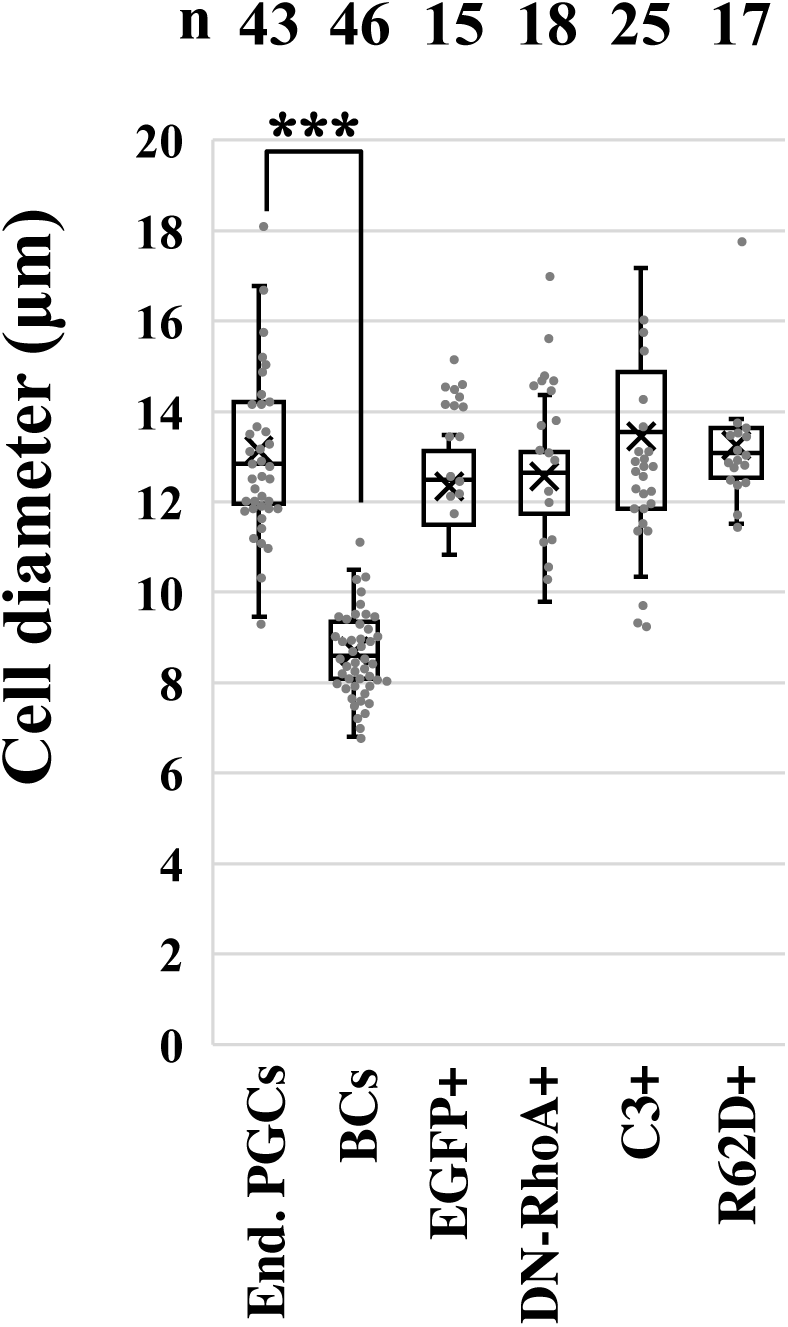
Cell diameters of endogenous PGCs and BCs, and manipulated cultured PGCs. Analyzed cell numbers are shown on top of the graph. ***p<0.001.

**Extended Data Figure 6.**
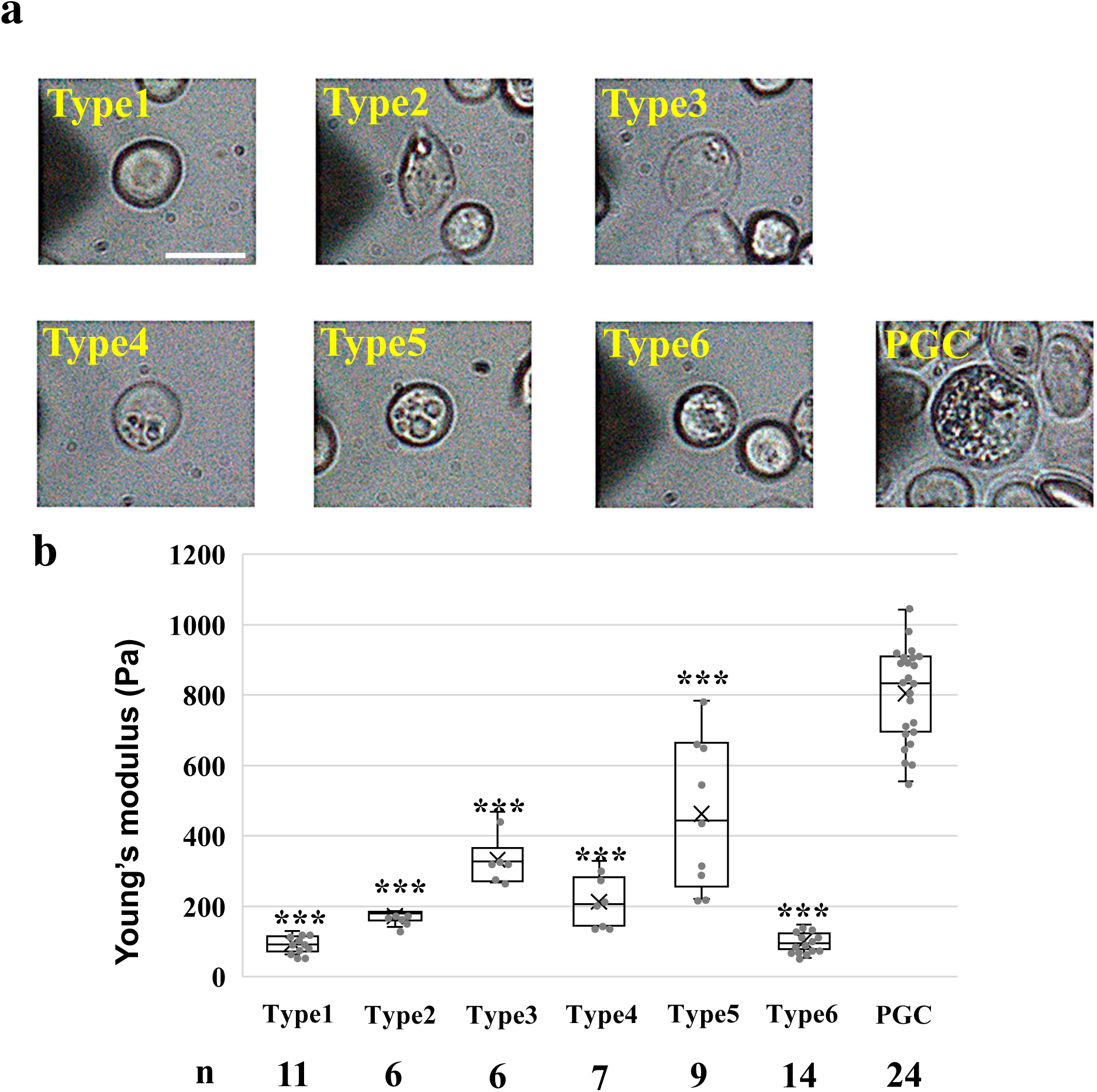
Seven types of intravascularly circulating cells subjected to the AFM indentation test to measure the cell stiffness. **a,** Appearance of cells harvested from the blood of HH15 chicken embryos. BCs are classified into 6 types (Type1 to Type6) by morphological differences. Scale bars: 10 μm. **b,** Quantification of cell stiffness of each cell type. Analyzed cell numbers are as indicated. ***p<0.001.

**Extended Data Figure 7.**
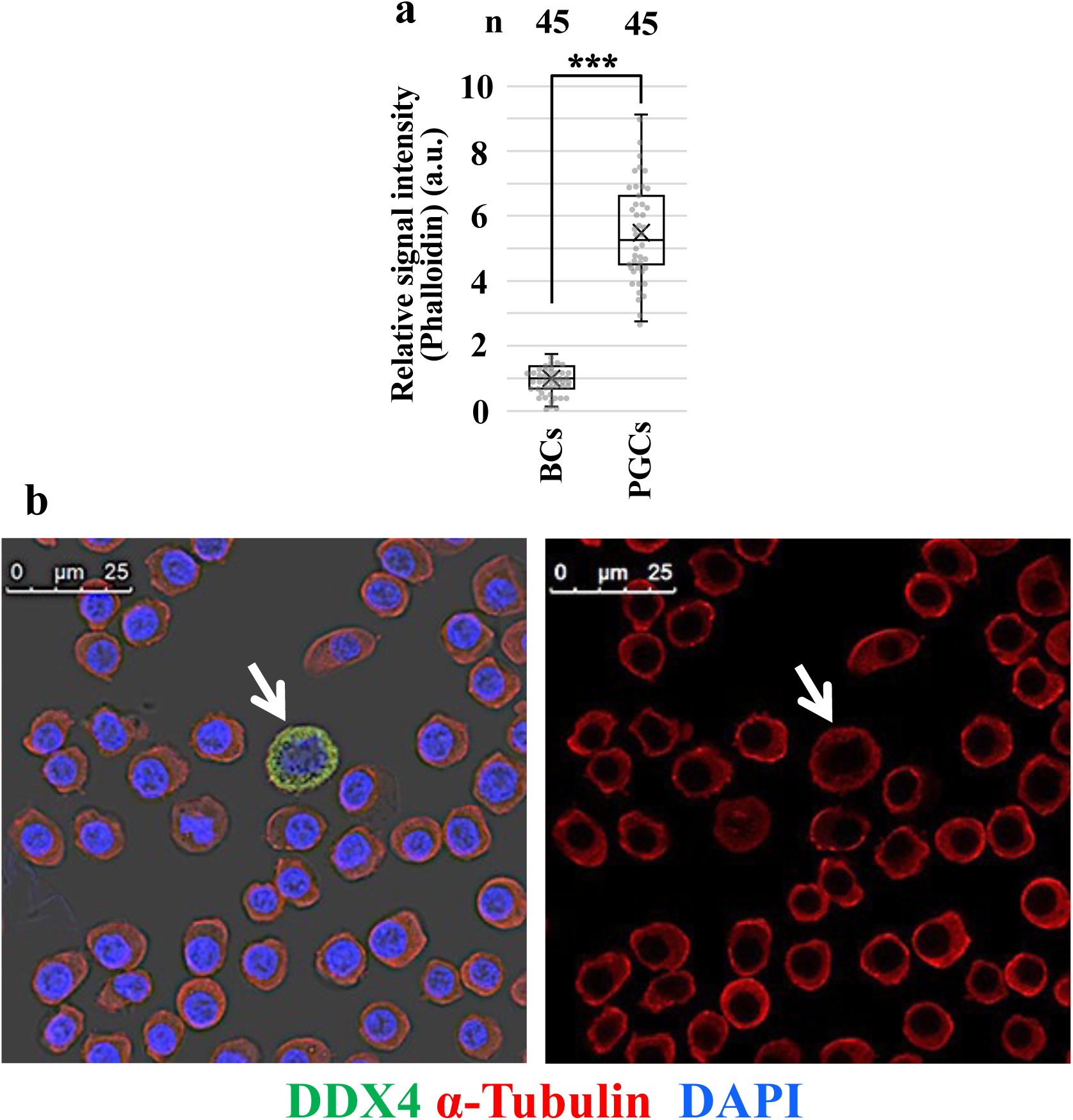
Signal intensity of phalloidin staining but not that of α-Tubulin is significantly greater in PGCs than BCs. **a,** Quantification of phalloidin staining signals in DDX4+ PGCs shown in Fig. 3c. Analyzed cell numbers are as indicated. ***p<0.001. **b,** Immunofluorescence of α-Tubulin (red) and DDX4 (green) with DAPI staining (blue) in BCs and PGCs (arrow) harvested from the blood of HH15 chicken embryos. Scale bars: 25 μm.

**Extended Data Figure 8.**
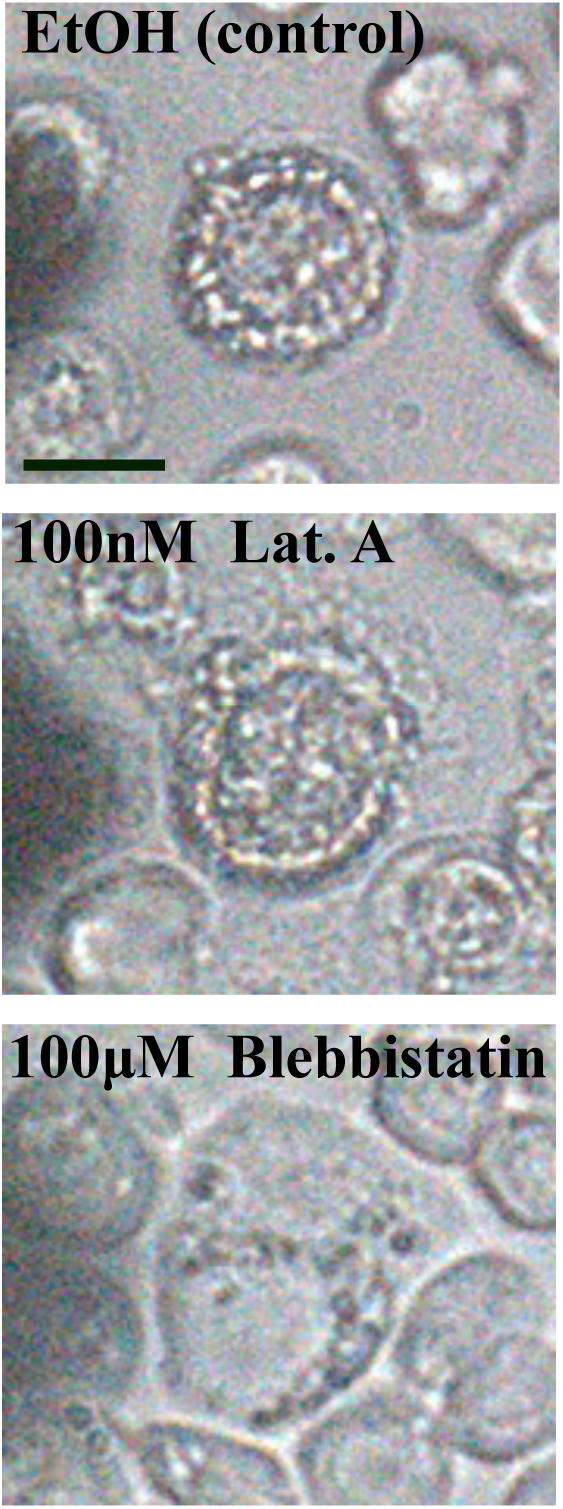
Appearance of endogenous PGCs treated by drugs. Appearance of PGCs treated by EtOH, 100nM Latrunculin A (Lat.A), or 100μM Blebbistatin. Scale bar: 10 μm.

**Extended Data Figure 9.**
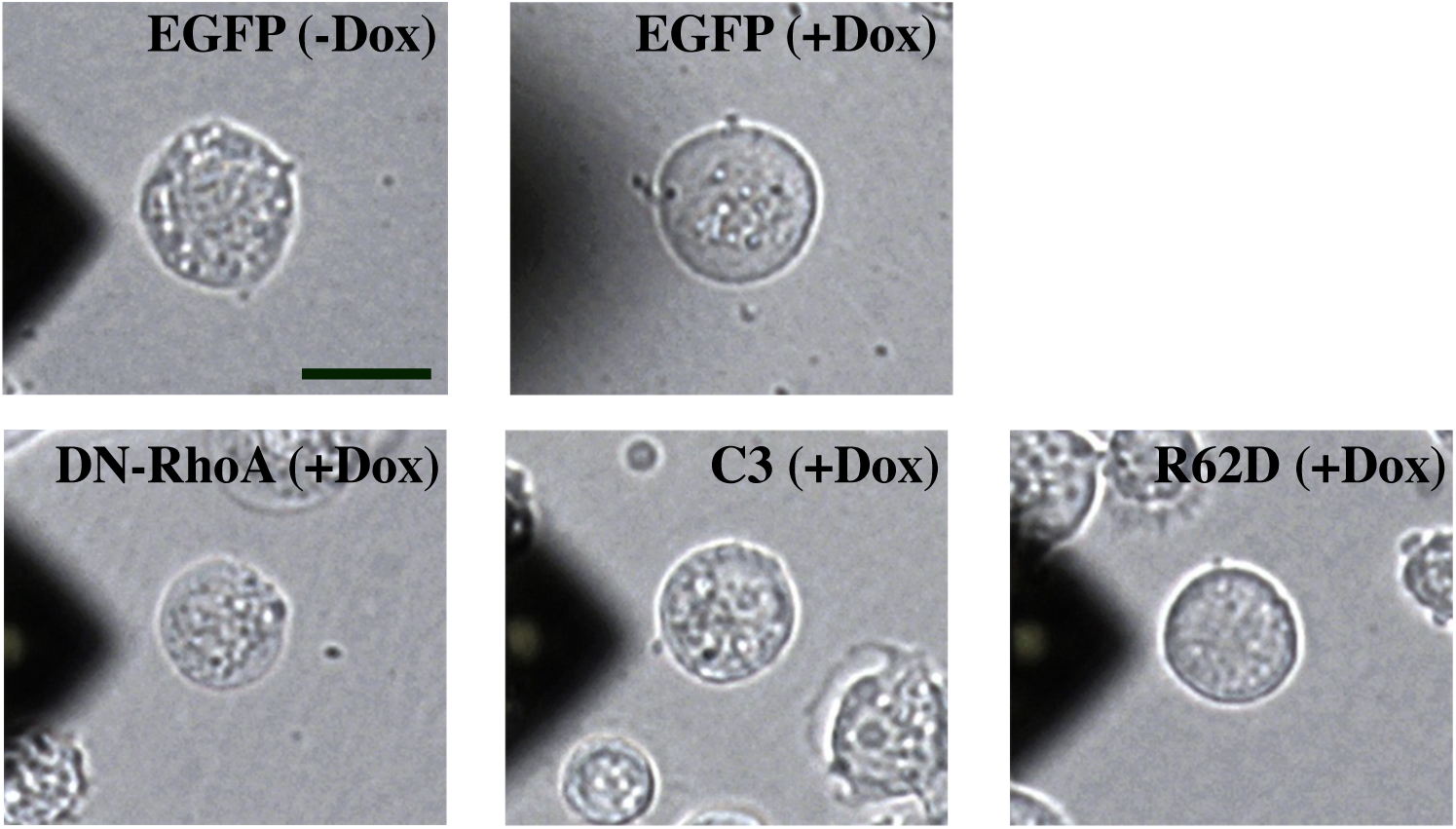
Appearance of gene-manipulated PGCs after Dox administration *in vitro*. Scale bar: 10 μm.

**Extended Data Figure 10.**
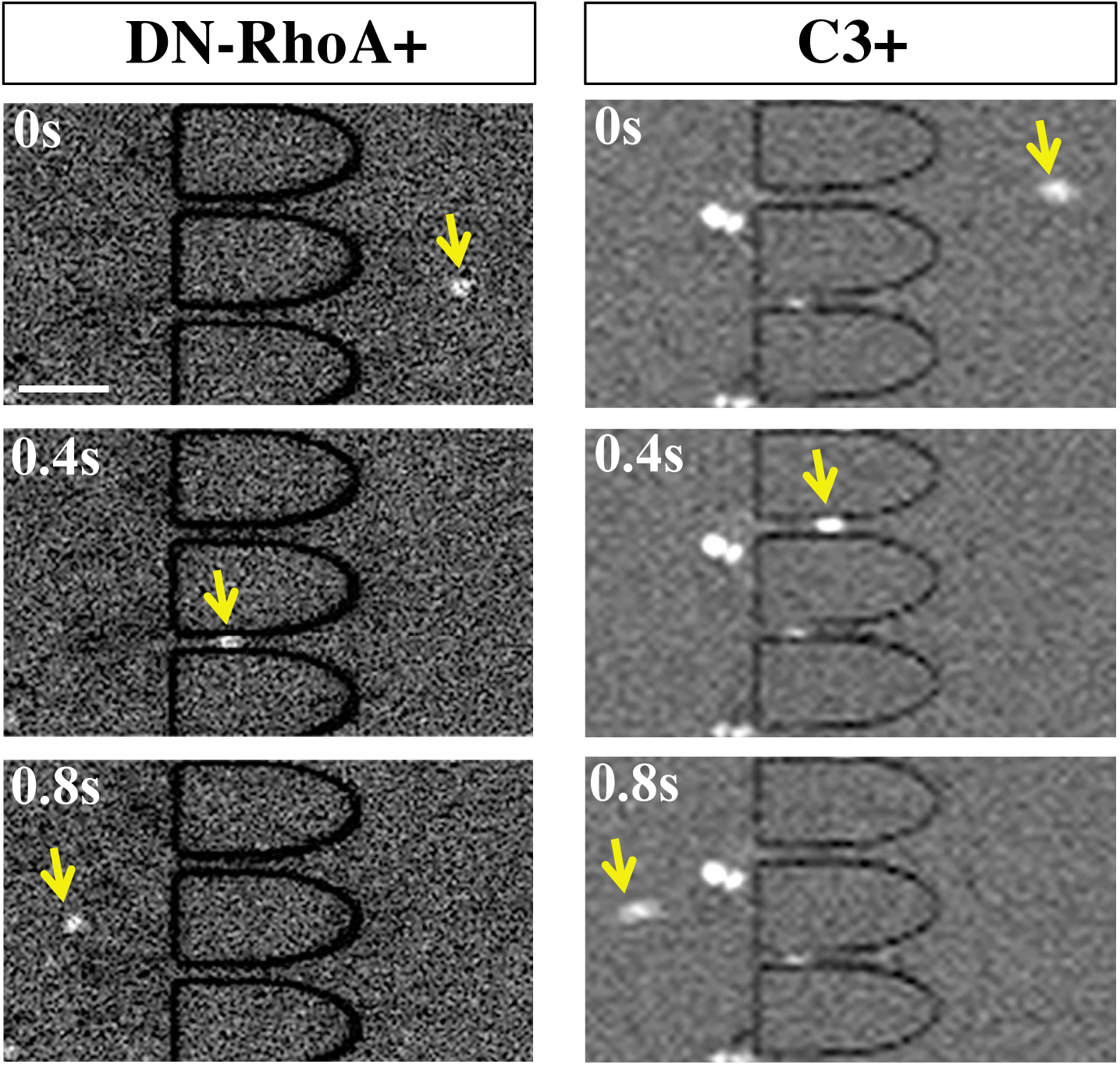
Microfluidic analyses of DN-RhoA- and C3-overexpressed PGCs. Motion captures from Supplementary videos 5 and 6. See legends of Figure 3g**, h.** Scale bar: 50 μm.

**Extended Data Figure 11:**
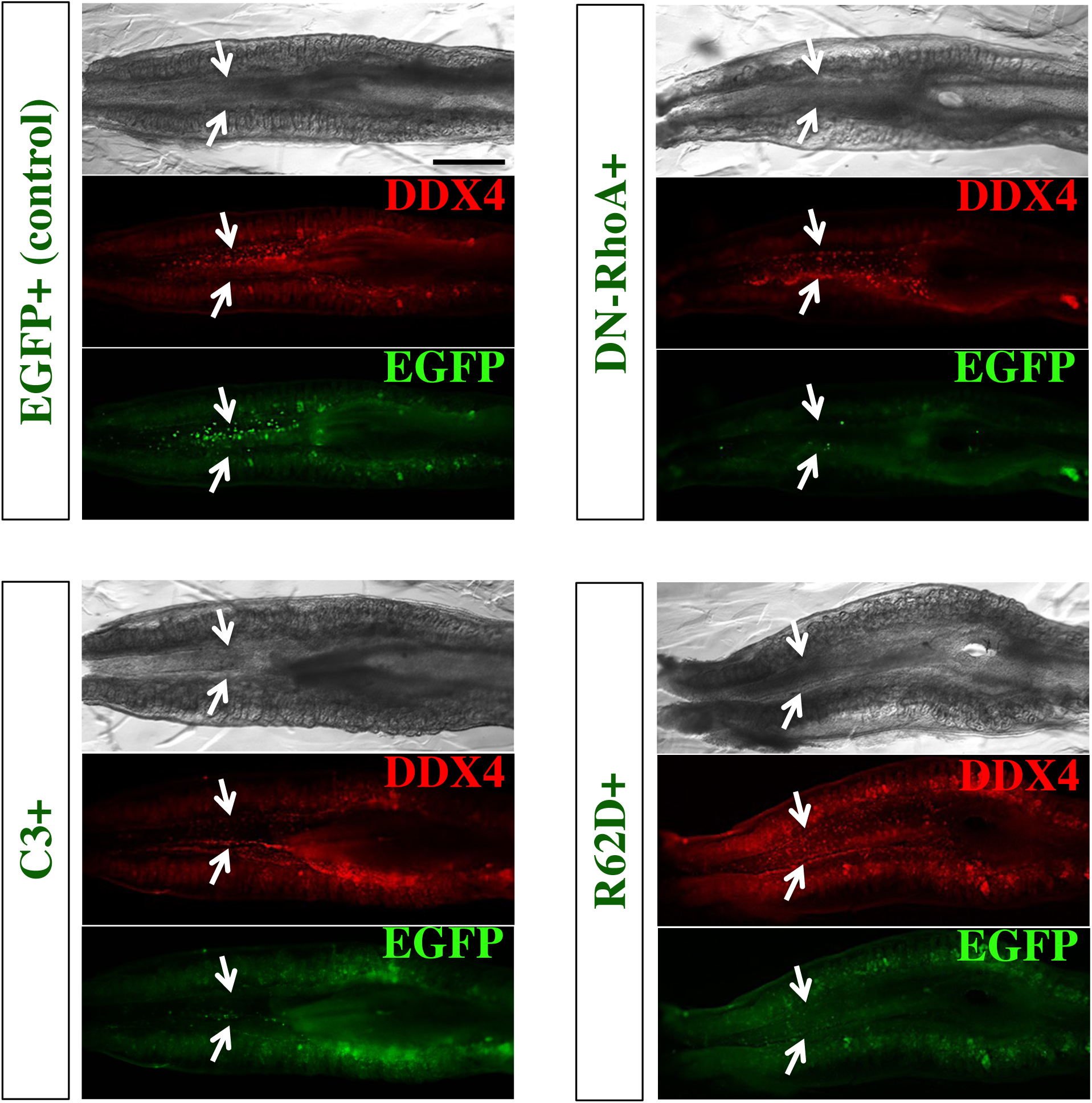
Localization of back-infused PGCs in the mesentery and forming gonads 2 days after infusion. Images of EGFP and DDX4 immunofluorescence in isolated tissues including mesonephros, gonads (white arrows), and dorsal mesentery located in between the gonads in E4.5 chicken embryos. EGFP+, DN-RhoA, C3+, or R62D+ PGCs were infused at HH15. Ventral view. Anterior is to the right. Scale bar: 500 μm.

**Extended Data Figure 12.**
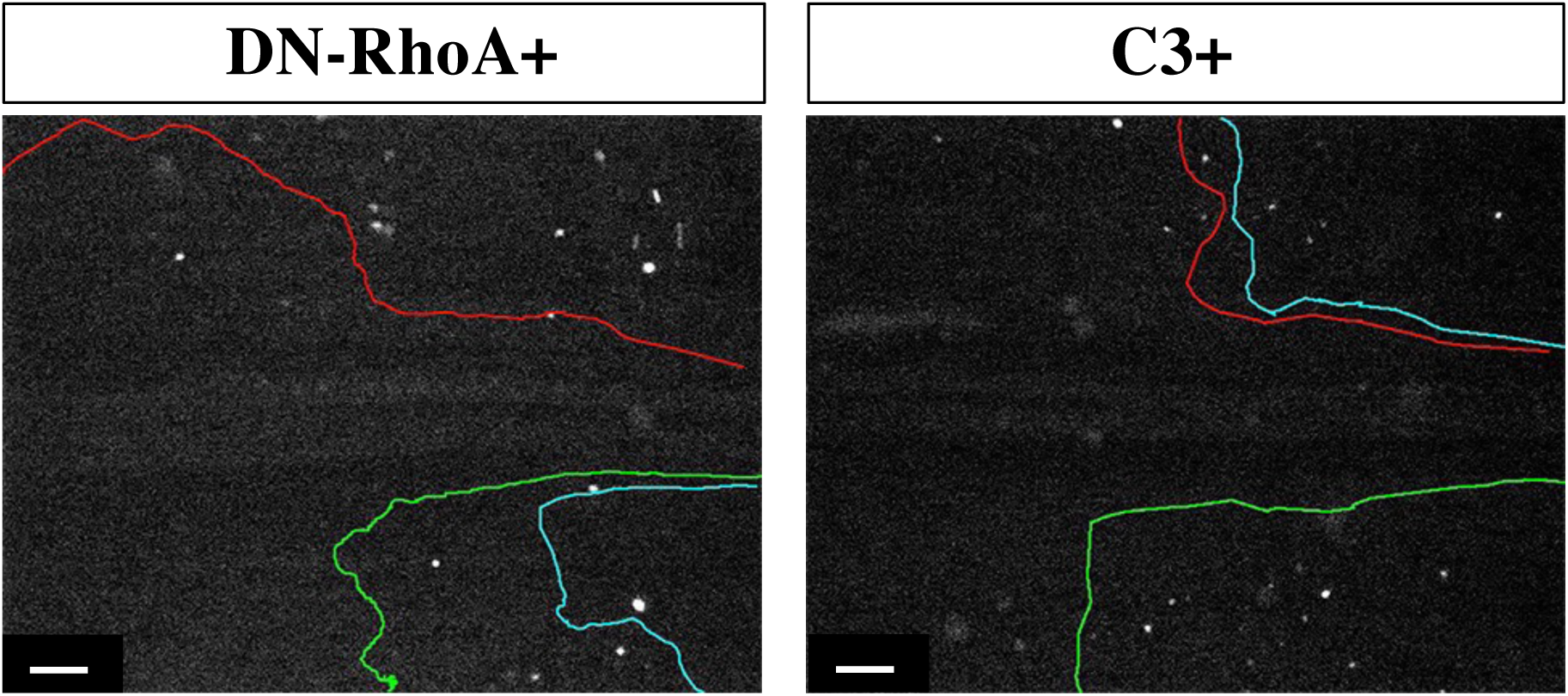
*In vivo* tracing of DN-RhoA- and C3-overexpressed PGCs in Ex-VaP. Movie captures taken from Supplementary videos 9 and 10. See legends of Figure 4a**, b.** Scale bars: 200 μm.

**Extended Data Figure 13.**
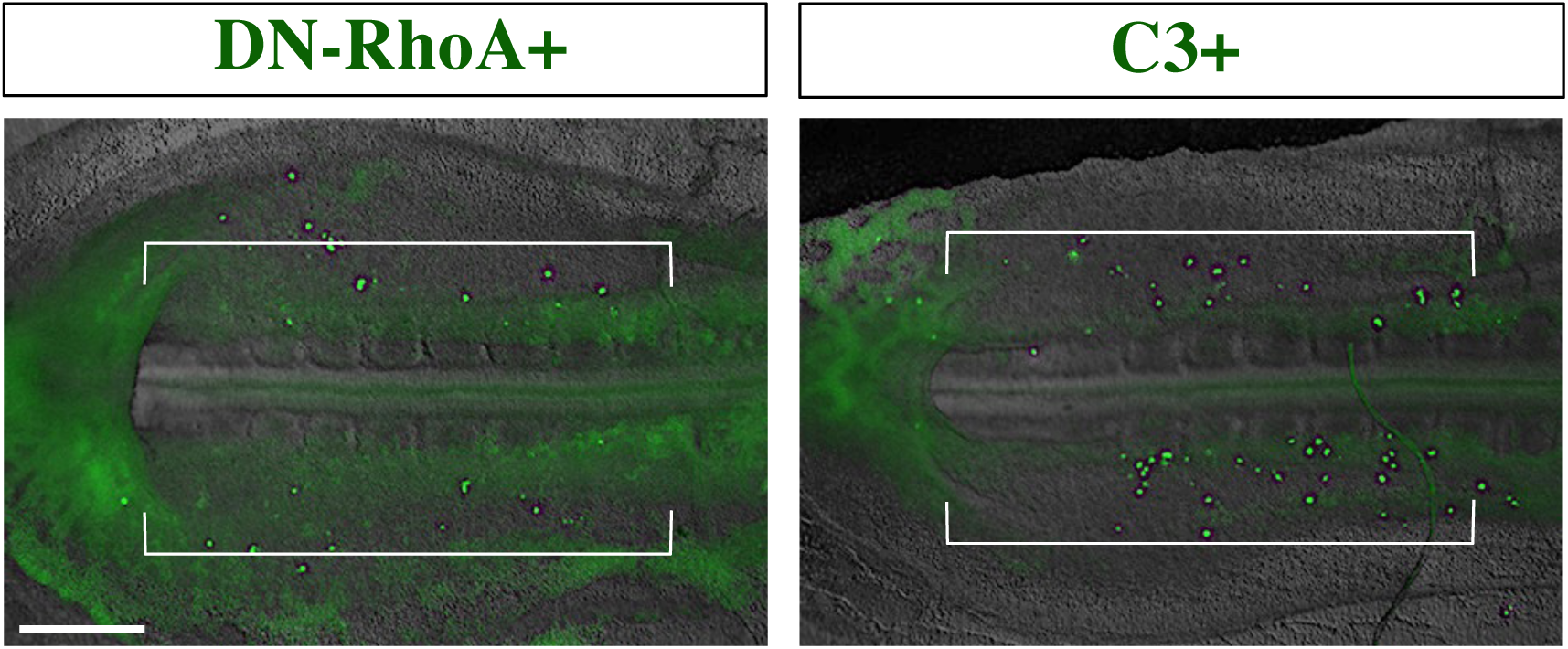
Localization in Ex-VaP of back-infused PGCs transfected with DN-RhoA- or C3- after 2 hours post-infusion. See legends of Figure 4d**, e.** Scale bar: 500 μm.

**Extended Data Figure 14.**
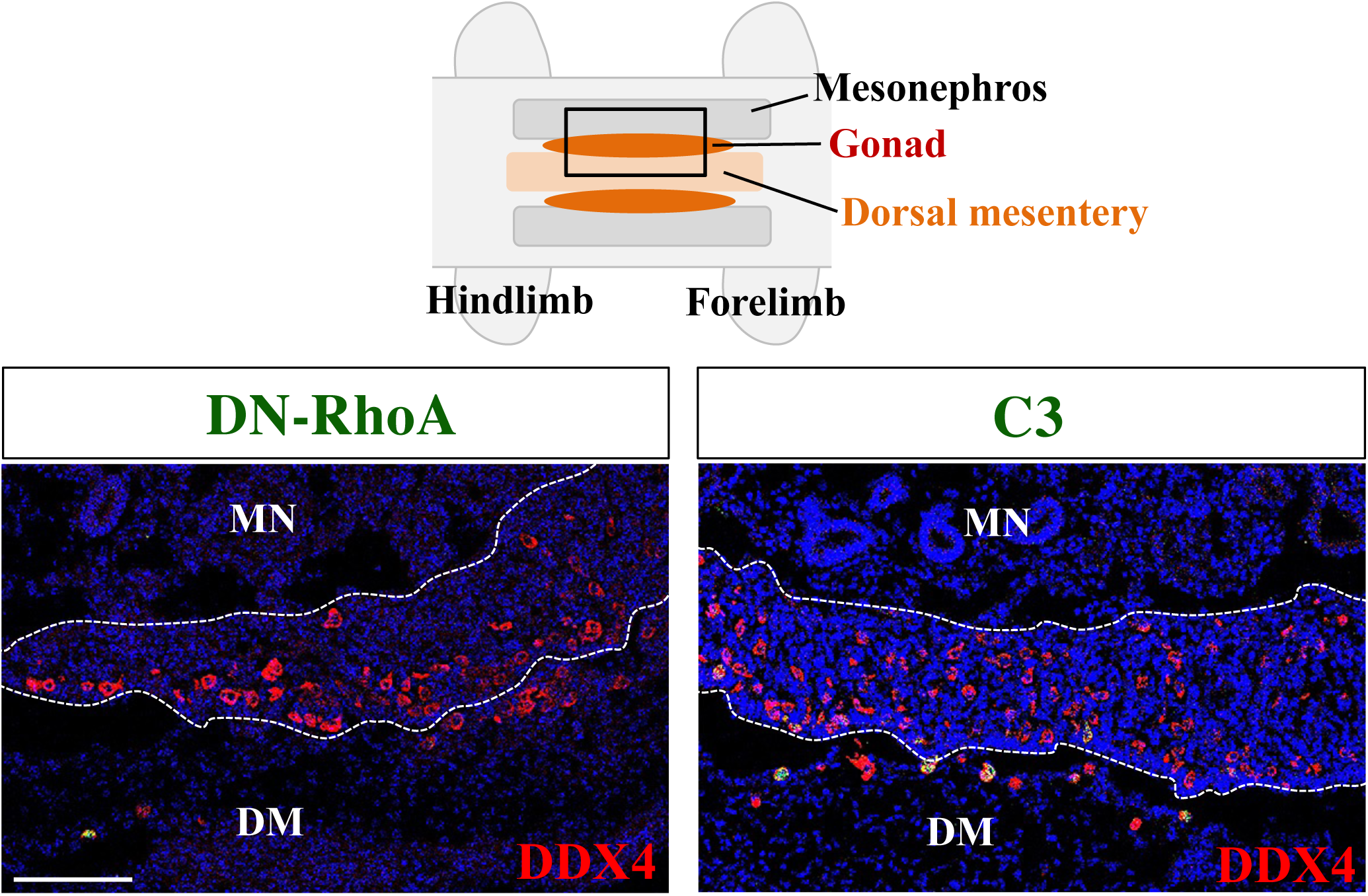
Cryo-sections of forming gonads and mesentery showing failed colonization of PGCs transfected with DN-RhoA- or C3 2 days post-infusion. The experiment are as explained in Fig 4g,h. Scale bar: 100 μm.

**Extended Data Figure 15.**
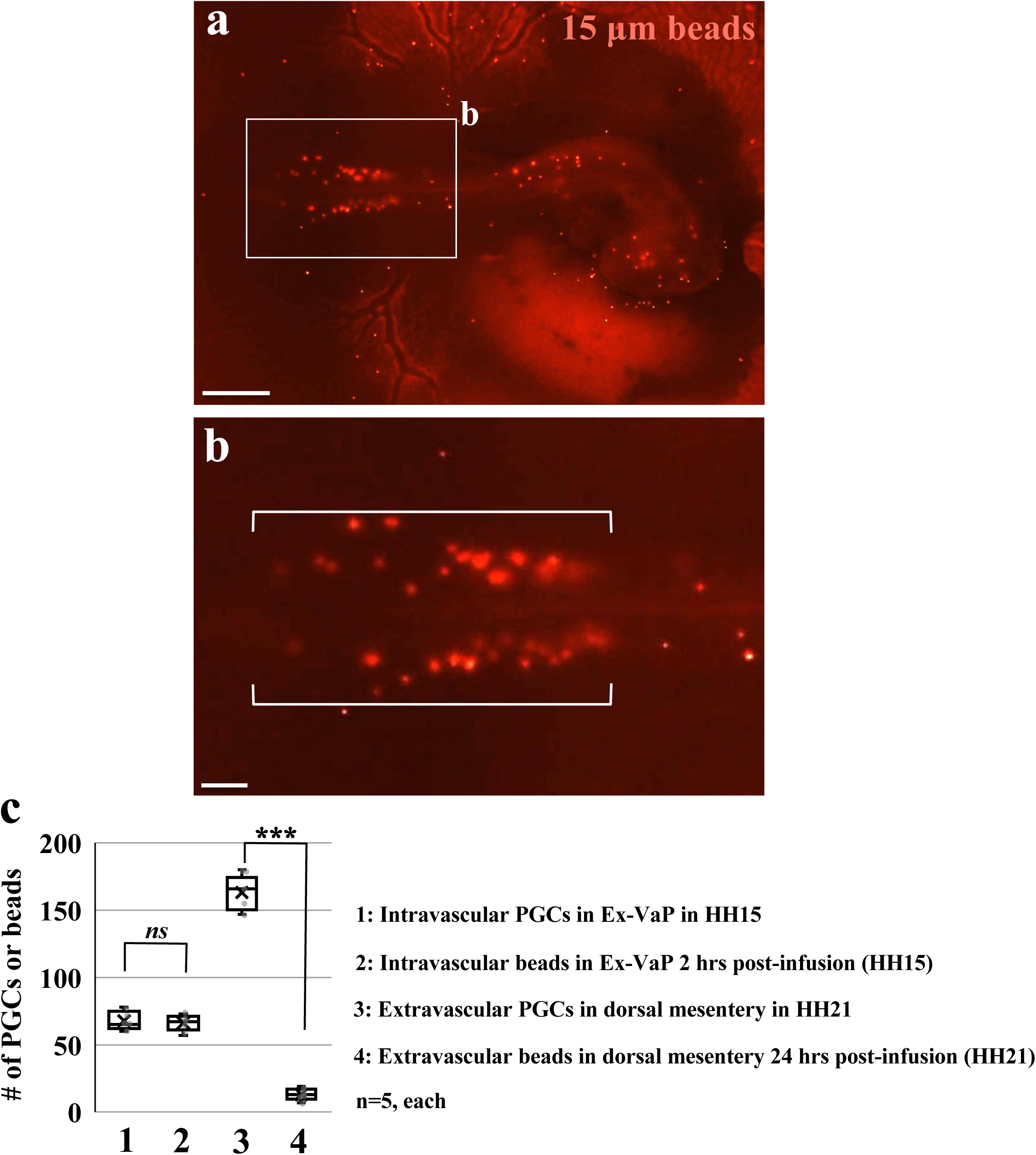
Localization and quantification of infused polystyrene beads of 15 μm diameter and endogenous PGCs in chicken embryos. **a,** Image of localization of 15 μm red fluorescent polystyrene beads in HH15 chick embryo. 2 hours after 1,000 beads were infused. **b,** Magnified image of the square in **a**. White brackets indicate the Ex-VaP area. **c,** Quantification of (1) intravascular endogenous PGCs in Ex-VaP at HH15, (2) intravascular beads in Ex-VaP 2 hrs after infusion at HH15, (3) extravascular endogenous PGCs in the dorsal mesentery at HH21, and (4) extravascular beads in the dorsal mesentery at HH21 24 hours post-infusion (n = 5, each). ***p<0.001. Scale bars: 1,000 μm in **a**, 250 μm in **b**.

